# Integrative genomic insights into parallel local adaptation to Danxia and Karst edaphic islands in the endangered tree *Firmiana danxiaensis*

**DOI:** 10.1101/2025.11.05.686652

**Authors:** Yali Li, Jinchu Luo, Lei Duan, Renchao Zhou, Jun Wen, Hongfeng Chen

**Affiliations:** Key Laboratory of National Forestry and Grassland Administration on Plant Conservation and Utilization in Southern China, South China Botanical Garden, Chinese Academy of Sciences, Guangzhou, Guangdong 510650, China; Guangdong Provincial Key Laboratory of Applied Botany, South China Botanical Garden, Chinese Academy of Sciences, Guangzhou, Guangdong 510650, China; University of Chinese Academy of Sciences, Beijing 100049, China; State Key Laboratory of Biocontrol and Guangdong Provincial Key Laboratory of Plant Stress Biology, School of Life Sciences, Sun Yat-sen University, Guangzhou, China; Department of Botany, National Museum of Natural History, Smithsonian Institution, Washington, DC, USA

**Keywords:** local adaptation, edaphic islands, Danxia landform, Karst landform, *Firmiana danxiaensis*, drought stress

## Abstract

Uncovering the genetic basis of local adaptation is a central goal in evolutionary biology. Edaphic islands, such as Karst (KL) and Danxia (DL) landforms, provide ideal natural laboratories for this endeavor. However, the mechanisms underlying parallel adaptation to contrasting edaphic extremes within a single species remain poorly understood. *Firmiana danxiaensis*, an endangered tree endemic to both KL and DL in the Nanling Mountains, offers a rare opportunity to address this question. We integrated a chromosome-level genome assembly, population genomic analysis of 225 individuals from 20 populations across its entire range, and genotype–environment association (GEA) with high-resolution soil and climate data. The relatively large genome of *F. danxiaensis* (1.51 Gb) reveals two ancient whole-genome duplications and a Quaternary burst of LTR retrotransposons, which, together with significant expanded drought-related transcription factor (TF) families, together constitute its genomic basis of adaptation. Population structure resolved four genetic lineages with distinct demography and inbreeding patterns. GEA identified stronger local adaptation in DL than KL, driven mainly by soil factors (e.g., cadmium, zinc), while climate variables (e.g., Bio9, Bio11) predominated in KL. Despite these divergent drivers, drought stress and elevated metal ion acted as major selective pressures in both landforms. Furthermore, six key TFs (e.g., *NAC090*, *bHLH42*, *MAPKKK20*) under both local adaptation and positive selection in DL are involved in drought response and metal ion binding, facilitating survival in stressful habitats. Our findings provide novel insights into parallel local adaptation to contrasting edaphic islands and offer concrete genomic guidance for the conservation of *F. danxiaensis*.

**Significance statement:** The genetic basis of parallel local adaptation to contrasting environments remains unclear. Using the endangered tree *Firmiana danxiaensis*, we uncover genomic mechanisms enabling its adaptation to both Karst and Danxia edaphic islands. We demonstrate that adaptation is primarily driven by soil factors in Danxia and climate in Karst, generating divergent genomic signatures yet converge on drought response. These findings establish a framework for understanding parallel local adaptation and offer concrete genomic guidance for conservation.

## Introduction

Uncovering the genetic basis of local adaptation is a central goal in evolutionary biology (Ostridge *et al*., 2025; Hoban *et al*., 2016; Postma & Agren, 2016; Bourgeois and Warren, 2021). Edaphic islands—spatially isolated soil habitats such as those on Serpentine, Karst, Danxia, Gypsum, and Dolomite (Palacio *et al*., 2007; Wang *et al*., 2017)—provide a powerful model for this endeavor. These habitats host edaphic specialists adapted to distinct soils, estimated 5–10% of all plant species (Corlett & Tomlinson, 2020; Hoorn *et al*., 2022; Wang *et al*., 2017; Rajakaruna, 2018), and thereby offer crucial insights into how genetic and environmental factors underpin survival under harsh and changing conditions (Allendorf *et al*., 2010; Ostridge *et al*., 2025).

However, the mechanisms underlying parallel local adaptation to contrasting edaphic extremes within a single species remain poorly understood. Previous research has advanced our understanding in two primary ways: by studying model or widespread species (Arnold *et al*., 2016; Wagner & Mitchell-Olds, 2018; Hahn *et al*., 2019; Liu *et al*., 2021), and by examining edaphic specialists restricted to a single soil type, such as Karst (Cao *et al*., 2023; Feng *et al*., 2020; Xie *et al*., 2021; Zhou *et al*., 2023; Jin *et al*., 2018) or Serpentine systems (Arnold *et al*., 2016; Sianta & Kay, 2019). Methodologically, studies have largely relied on reciprocal transplants, common gardens, and selective sweep scans (Pavlidis *et al*., 2013; Hu *et al*., 2021; VanWallendael *et al*., 2019), with limited use of of genotype-environment association (GEA) analyses—a powerful approach for detecting loci associated with environmental variables (Abebe *et al*., 2015; Hohenlohe *et al*., 2021). Furthermore, even when GEA is applied, most studies prioritize climatic variables over soil parameters, despite evidence that soil properties exert strong, fine-scale selection in these systems (Baxter & Dilkes, 2012; Wang *et al*., 2017; Sang *et al*., 2022; Marková *et al*., 2023; Lazic *et al*., 2024). Practical difficulties in soil sampling may underlie this bias. Consequently, the genetic basis of local adaptation in edaphic islands remains poorly resolved, including the relative roles of soil versus climate, the identity of primary selective agents, and the nature of underlying adaptive genes.

The Nanling Mountains, the largest east–west trending range in subtropical East Asia, form a major biodiversity hotspot (Lu *et al*., 2023; Zhao *et al*., 2024). This region harbors two contrasting edaphic islands—Danxia (DL) and Karst (KL) landforms—which support exceptional levels of endemism and are recognized both as UNESCO World Natural Heritage Sites and IUCN global centres of plant diversity (Davis & Heywood, 1997; Wang *et al*., 2017), making them ideal models for studying parallel local adaptation in edaphic specialists. Both landforms feature shallow, erosion-prone, drought-stressed, and metal-rich soils (Hao *et al*., 2015), yet differ fundamentally in chemistry: DL soils derived from red sandstone are acidic and nutrient-poor (Hua, 2000), while KL soils are calcium-rich, slightly alkaline, and high in magnesium (Hao *et al*., 2015). This striking environmental heterogeneity imposes distinct selective pressures expected to drive divergent local adaptation (Wang *et al*., 2025; Savolainen *et al*., 2013).

The present study focuses on *Firmiana danxiaensis*, a critically endangered tree species of the cotton family (Malvaceae) endemic to the Nanling Mountains (Qin *et al*., 2017) with high ornamental and ecological value. Unlike most edaphic specialists confined to a single soil type, *F. danxiaensis* exhibits a unique distribution across both DL and KL. The occurrence of this species on both landforms thus provides a rare opportunity to uncover the genetic mechanisms underlying parallel local adaptation to divergent soil environments. Moreover, elucidating such local adaptation is crucial for informing conservation strategies under ongoing climate change (Armstrong *et al*., 2021; Ostridge *et al*., 2025). Local adaptation results from the balance between selection and other evolutionary forces—such as gene flow, effective population size, and historical demography (Hoban *et al*., 2016; Ostridge *et al*., 2025). Therefore, assessing these population genetic features is essential for guiding effective conservation management and genetic rescue efforts (Bourgeois & Warren, 2021; Fuentes-Pardo & Ruzzante, 2017).

To address these questions, we first generated a chromosome-level de novo assembly of the whole genome of *F. danxiaensis* and resequenced 225 individuals from 20 populations spanning its natural distribution. By integrating high-resolution data on field measurements of 20 soil variables and climatic data with comparative genomics, population genomic, and GEA approaches, we aim to: (1) characterize the genomic features and adaptive evolutionary history of *F. danxiaensis*; (2) quantify how evolutionary forces, such as gene flow and demographic history shape population differentiation; (3) dissect the genetic basis of parallel local adaptation to Danxia and Karst landforms and evaluate the relative contributions of soil and climate factors; and (4) delineate conservation units based on genomic and environmental insights to guide targeted conservation strategies. This study advances our understanding of genetic mechanisms underlying parallel local adaptation in distinct edaphic islands and provide an integrative framework for future research on other edaphic specialists.

## Results

### Chromosome-scale genome assembly and annotation of *Firmiana danxiaensis*

Genome survey using *K*-mer analysis of Illumina reads revealed that the genome size of *F. danxiaensis* was about 1.53 Gb (Figure S1), which was consistent with the results of flow cytometry analysis (Figure S2; Table S1). We generated a genome assembly of 1.51 Gb, with contig N50 of 69.08 Mb and 97.37% of the contig sequences were anchored on 20 pseudo-chromosomes (Figure 1a, b; Table S2). The BUSCO analysis further demonstrated that 1,594 of the 1,614 core genes were complete, showing relatively high completeness (98.8%) of the assembled genome (Table S3). We predicted a total of 31,938 protein-coding genes (Table S4), in which 31,218 (97.75%) were functionally annotated (Table S5). The genome of *F. danxiaensis* contained a total of 1.01 Gb of repetitive sequences, accounting for 83.12% of its entirety, with LTR-RTs being the most abundant at 67.12%. Among the LTR-RTs, Copia and Gypsy were the most common superfamilies, representing 2.52% and 25.16% of genome, respectively (Table S6). Non-coding RNAs information can be found in Table S7. Taken together, we generated a high-quality, chromosome-scale genome of *F. danxiaensis*.

**Figure 1.**
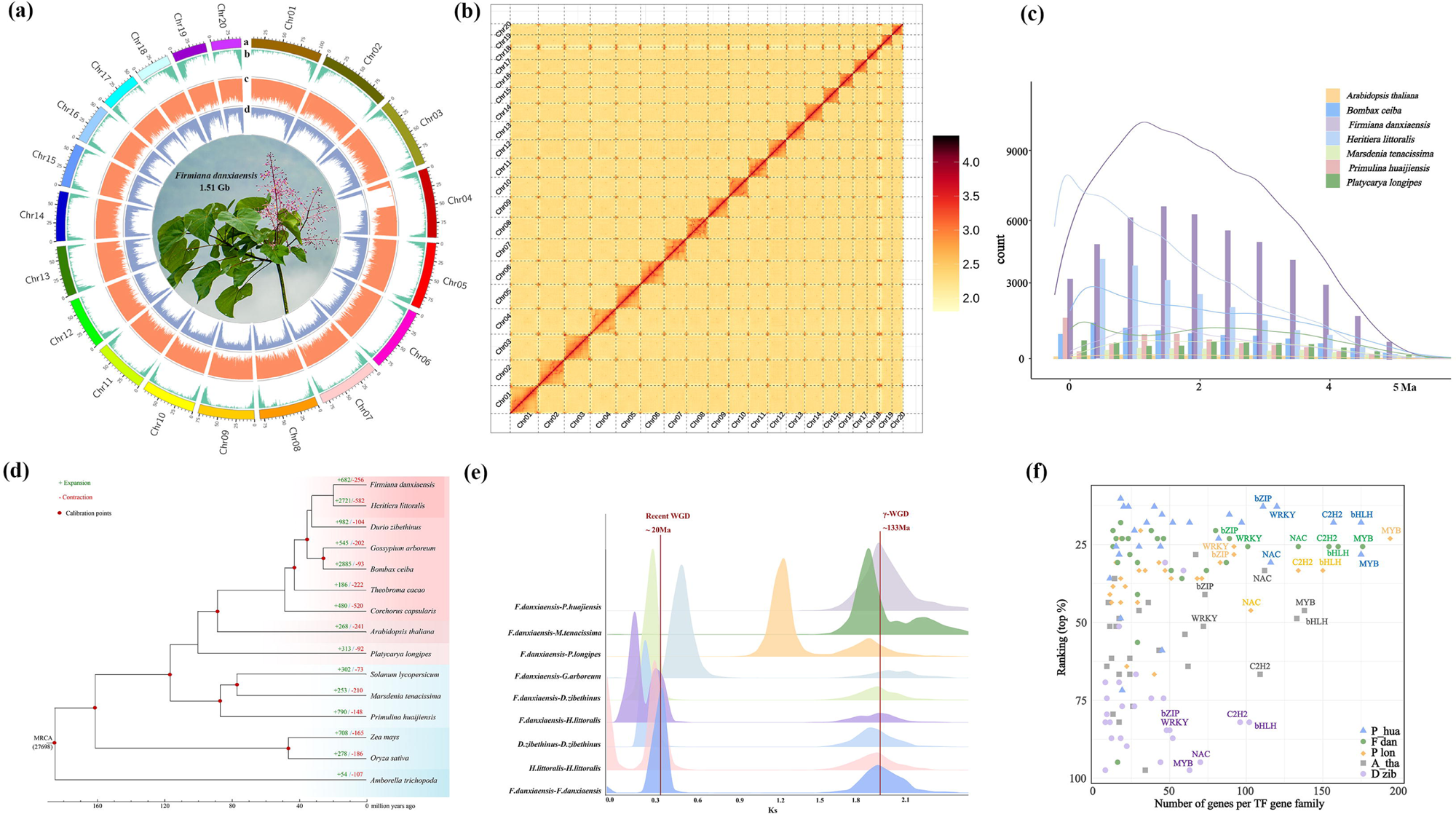
Whole genomic evidence of adaptation in *Firmiana danxiaensis.* (a) Circos plot showing genomic features of the assembled genome. Tracks from outermost to innermost represent chromosome positions a), gene density b), SNP density c), and indel density d). (b) Hi-C contact map of the *F. danxiaensis* genome. (c) Insertion times of LTR retrotransposons (LTR–RTs) in seven species. (d) Phylogenetic tree of 15 species based on concatenated alignments of 331 single-copy orthologous genes. Estimated divergence times and the corresponding time scale are shown at the bottom. The red solid circles indicate the calibration nodes. (e) Distribution of synonymous substitution rates (Ks) among paralogous and orthologous gene pairs in *F. danxiaensis*, *Primulina huaijiensis*, *Marsdenia tenacissima*, *Platycarya longipes*, *Gossypium arboreum*, *Durio zibethinus*, and *Heritiera littoralis*. (f) Analysis of transcription factor (TF) families in *F. danxiaensis*, *P. huaijiensis*, *P. longipes*, *D. zibethinus*, and *Arabidopsis thaliana*. The x-axis shows the number of TFs in each species. the y-axis indicates the rank of each TF family compared to 35 other eudicot species.

### Genome-scale evidences that may facilitate adaptation

The activity of TEs not only influences genome size but can also induce various genetic variations, thereby driving lineage-specific diversification and adaptation(Oliver *et al*., 2013; Warren *et al*., 2015). To investigate whether TE activity contributed to the adaptation of *F. danxiaensis*, we analyzed the insertion history of LTR-RTs. Our results indicated that LTR insertion was a continuous process, with approximately 40% of the insertions occurred 1∼2.5 Ma (Figure 1c). A comparison with six other plant species highlighted notable variations in both the number of LTR-RT insertions and their peak periods. *F. danxiaensis* exhibited the highest number of insertions, peaking at 1.3∼1.4 Ma, which distinguished from its relatives *H. littoralis* and *B. ceiba* (Figure 1c; Table S8). These results indicate that rapid accumulation of LTR-RT insertions may have played a crucial role in the adaptation of *F. danxiaensis* to the climatic condition during the quaternary glaciation.

Whole-genome duplication (WGD) is a widespread phenomenon in plants and is widely regarded as a major force driving genomic evolution, accelerating speciation, and promoting adaptive divergence(Soltis *et al*., 2014). To trace the evolutionary trajectory of WGD events in *F. danxiaensis*, we identified and dated them in its genome. The analysis showed that *F. danxiaensis* and *Heritiera littoralis*, the closest relative with a sequenced genome, shared two WGD events, which occurred approximately 133 and 20 million years ago (Ma) (Figure 1e). The two WGD events aligned with the γWGD and the αWGD events, which were observed in many other angiosperms (Jiao et al. 2011; Zhang et al. 2020). Their timing suggests a potential link to major episodes of climatic and environmental change, possibly contributing to adaptive evolution in the *F. danxiaensis*. Compared to the most recent common ancestor of *F. danxiaensis* and *H. littoralis*, the genome of *F. danxiaensis* has undergone significant expansion in 682 gene families (Figure 1d), with gene ontology enrichment analysis revealing a strong association with hydrolase activity, nucleic acid binding, and metal ion binding (Figure S3). Notably, the expansion of metal ion binding-related functions may represent a key genomic adaptation to the metal-rich soils characteristic of both the DL and KL.

Additionally, in eudicots, transcription factors (TFs) form large superfamilies, with duplicated genes playing key roles in adaptive evolution (Lehti-Shiu & Shiu, 2012). To investigate the potential TFs contributing to the adaptation of *F. danxiaensis,* a total of 2,238 TFs were identified in the *F. danxiaensis* genome, taking up 7.00% of all genes (Table S9). TFs from 36 eudicots were calculated and compared with those of *F. danxiaensis* (Feng *et al*., 2020) (Table S10). Relative to two generalist species (*Arabidopsis thaliana*, *Durio zibethinus*), *F. danxiaensis* and two KL endemic specialists (*Primulina huaijiensis*, *Platycarya longipes*) had significantly more members in six drought-related TFs families (*bHLH*, *MYB*, *C2H2*, *NAC*, *bZIP*, and *WRKY*) (Hao *et al*., 2021; Shinozaki & Yamaguchi-Shinozaki, 2007; Singh *et al*., 2002; Wu *et al*., 2009) (Figure 1f). Furthermore, *bHLH*, *WRKY*, *bZIP, TCP,* and *DREB* families exhibited significant expansion (p < 0.01) and were actively involved in the local adaptation and positive selection of *F. danxiaensis* to both DL and KL (Table S11). These results collectively indicate that the expansion of these TF families likely plays a critical role in the adaptation of *F. danxiaensis* to drought-prone environments in DL and KL.

### Phylogeny, population structure, genetic diversity and inbreeding dynamics

We resequenced 225 individuals of *F. danxiaensis* from 20 wild populations at all six known sites (Figure 2b). The average sequencing depth of these individuals was 10.02×, covering 99.55% of the genome on average (Table S12). Single-nucleotide polymorphism (SNP) calling following strict quality control criteria resulted in a set of 5,471,157 high-quality SNPs that were used for following analyses. Population structure of *F. danxiaensis* aligned well with its geographic distribution and landform types, individuals in close proximity tend to lineage together (Figure 2c). The phylogenetic tree based on SNPs (Figure 2d) and PCA results (Figure 2e) also supported four distinct lineages within *F. danxiaensis* (Figure S4). Structure analysis showed that populations from the Baiwan (BW) and Yingde (YD), inhabiting in KL (Figure 2a, b), formed one lineage (BWYD). Those from Shixing (SX) and Nanxiong (NX) in DL formed another lineage (SXNX) (Figure 2a, b), while Danxia (DX) and Baizhangyan (BZY), also in DL, each formed independent lineages.

**Figure 2.**
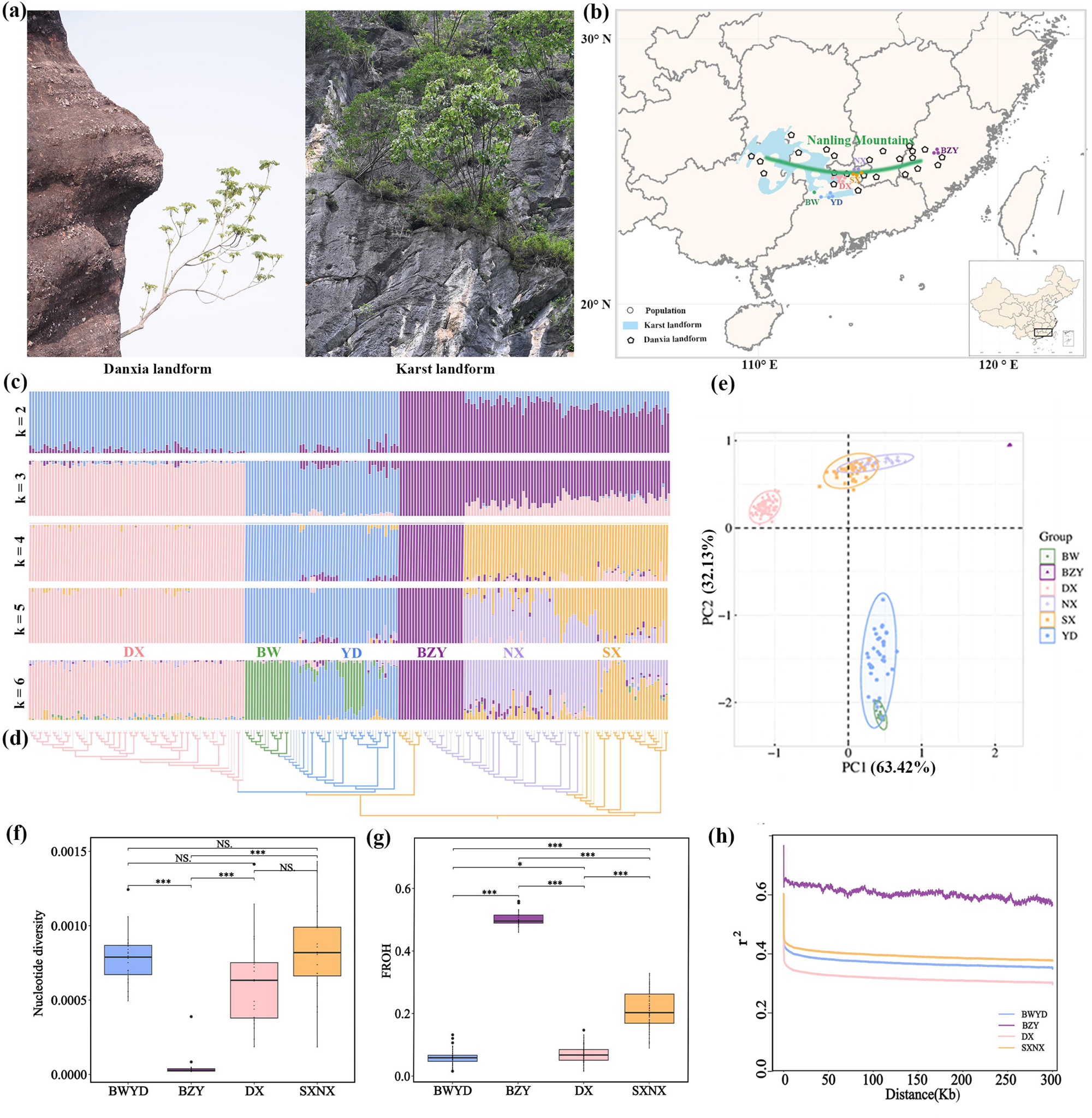
Habitat, population structure, and genetic diversity of *Firmiana danxiaensis.* (a) Representative habitats of *F. danxiaensis* in Danxia and Karst landforms. (b) Sampling across 20 wild populations. Solid circle represents sampled population of *F. danxiaensis*. Karst landform: BW and YD sites, and Danxia landform: BZY, DX, SX, and NX sites. Green line represents the ridgeline of Nanling Mountains. The blue band-shaped area represents karst landforms, and the five-pointed star shape represents Danxia landforms. (c) Population structure inferred using ADMIXTURE. Each bar on the x-axis represents an individual; the y-axis indicates ancestry proportions. (d) Maximum-likelihood phylogeny reconstructed with IQ-TREE. Branches with bootstrap support values ≥ 90 were highlighted in bold. (e) Principal components analysis (PCA) of 225 individuals of *F. danxiaensis*. (f) Nucleotide diversity in four genetic lineages (π) of *F. danxiaensis*. (g) Proportion of runs of homozygosity (ROH) relative to total genome length (FROH). (h) Linkage disequilibrium (LD) decay.

For whole-genome population statistics, pairwise fixation index (*F*_ST_) among four lineages ranged from 0.41 to 0.75, indicating extremely high genetic differentiation(Wright, 1984) (Table S13). Nucleotide diversity (π) varied across clusters: BZY (4.94 × 10^-5^) < DX (7.27 × 10^-4^) < BWYD (7.78 × 10^-4^) < SXNX (8.21 × 10^-4^) (Figure 2f). To further assess the inbreeding level of *F. danxiaensis*, we calculated the individual inbreeding coefficient (*F*_IS_) for all 225 samples. The average individual inbreeding coefficient (*F*_IS_) values across the four lineages were negatively correlated to π: BZY (0.96) > DX (0.59) > BWYD (0.54) > SXNX (0.46) (Figure S5a). Historical population dynamics and recent inbreeding can be detected through runs of homozygosity (ROH) in the genome (Kardos *et al*., 2018). We identified ROH longer than 500 kb in all *F. danxiaensis* individuals (Figure S5b) and calculated the proportion of ROH length to the total genome length (FROH) for each individual. BZY showed the highest average *F*_ROH_ (0.50), followed by SXNX (0.21), DX (0.07), and BWYD (0.06) (Figure 2g), highlighting that BZY has experienced particularly severe and recent inbreeding event. Moreover, whole-genome heterozygosity (het) (Figure S5c) was significantly negatively correlated with *F*_ROH_ (Figure S5d). The recombination rate (Figure S5e) and linkage disequilibrium decay patterns (Figure 2h) further supported these findings.

### Demographic history and gene flow

To evaluate the effective population size (Ne) trajectories of the four *F. danxiaensis* lineages, we applied Fitcoal2 (Hu *et al*., 2023) based on the site frequency spectrum (Figure 3a). The results revealed distinct yet climatically influenced demographic histories. The BZY lineage maintained a consistently small Ne since the Middle Pleistocene (∼0.70 Ma) until a pronounced expansion in the early Holocene (∼0.011 Ma). In contrast, the BWYD lineage experienced a sharp expansion during the Middle Pleistocene (∼0.64 Ma), remained stable for an extended period, and then entered continuous contraction from the Late Pleistocene (∼0.083 Ma). The SXNX lineage expanded steadily through the Middle Pleistocene, underwent a brief but marked expansion during the Penultimate Glaciation (PG, 0.13∼0.30 Ma) (Zheng *et al*., 2002), and subsequently declined. Similarly, the DX lineage grew exponentially from the Middle Pleistocene (∼0.55 Ma), peaked around ∼0.25 Ma, and sharply contracted during the PG. Notably, all lineages except BZY experienced a population bottleneck during the Last Glacial Maximum (LGM, 0.024∼0.011 Ma) (Zheng *et al*., 2002). These patterns were further supported by Fastsimcoal2 simulations (Excoffier *et al*., 2013), which modeled 30 demographic scenarios with and without gene flow; the best-supported model based on AIC values was consistent with the Fitcoal2 inferences (Figure 3b; Table S14).

**Figure 3.**
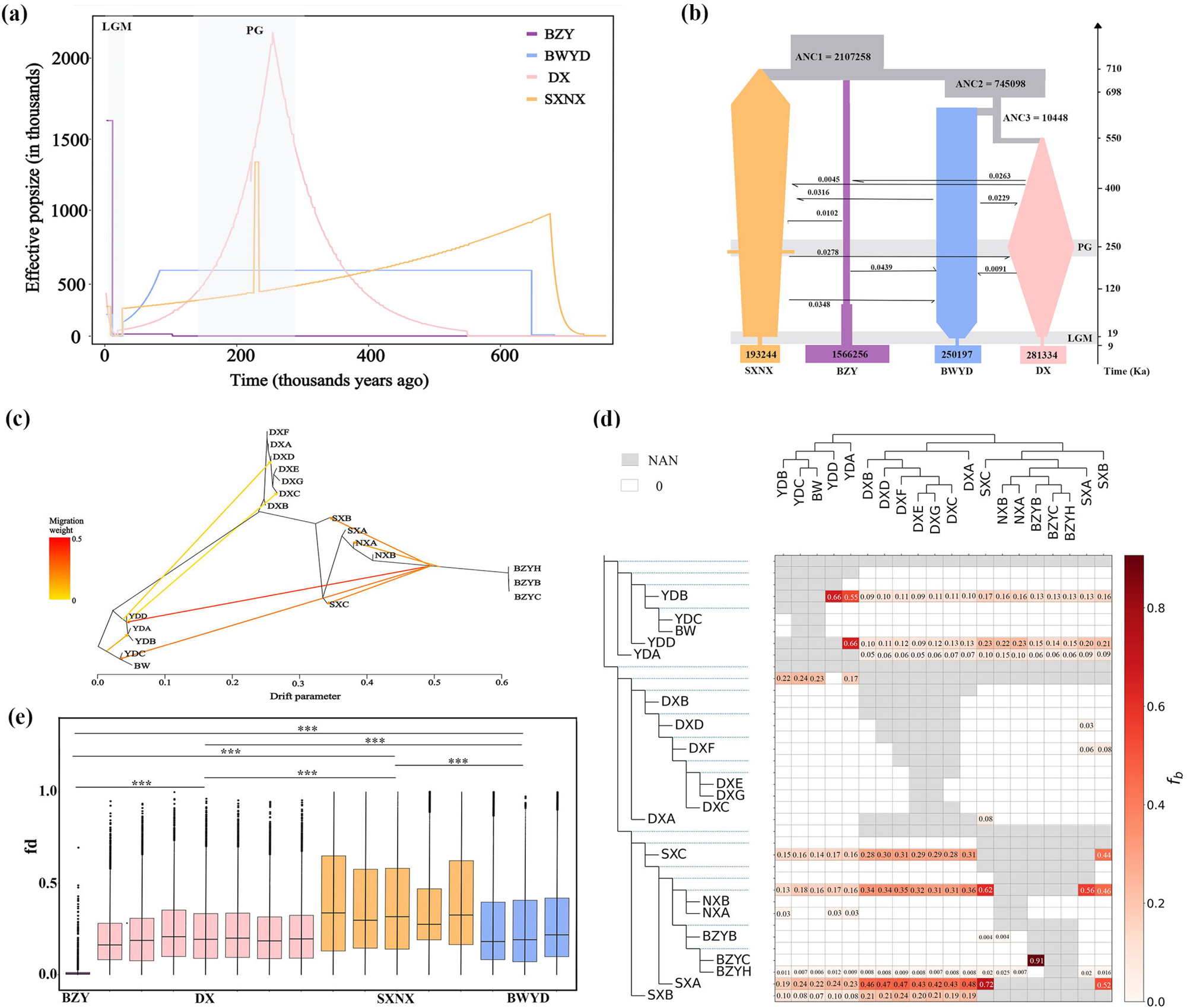
Demographic history and gene flow of *Firmiana danxiaensis.* (a) Changes in effective population size (Ne) inferred by FitCoal. (b) Demographic history inferred by Fastsimcoal2. (a,b) BZY, DX, SXNX, and BWYD represent four lineages of *F. danxiaensis*. (c) Maximum-likelihood tree and inferred migration events among populations. Arrows indicate gene flow direction, with darker red representing higher likelihood. (d) f-branch values among *F. danxiaensis* populations. Gray squares = missing; white = f-branch value of 0; red = presence of f-branch values (darker = higher); numbers indicate f-branch value. (e) The fd value quantifies the average proportion of gene flow in the genome, calculated using 10 kb windows.

In addition, to investigate gene flow among four lineages of *F. danxiaensis*, we integrated two complementary methods. Treemix (Pickrell & Pritchard, 2012) enables the analysis of gene flow direction. The strongest gene flow occurred from SXNX branch to BWYD (Figure 3c). To further quantify the gene flow, we applied Dsuite (Malinsky *et al*., 2021). The ABBA-BABA test heatmap showed that SXNX exhibits a higher level of gene flow (Figure S6). Additionally, fb values were calculated to assess gene flow within specific branches. The findings revealed that gene flow within *F. danxiaensis* populations is higher than between populations, with the strongest inter-population gene flow occurring between the SXNX and DX lineages (Figure 3d). Furthermore, the fd value of Dinvestigate model results indicated that SXNX experiences the highest level of gene flow, while the BZY was the least affected (Figure 3e). Overall, the results from these complementary approaches reveal a spectrum of gene flow among the lineages, with SXNX exhibiting the highest level and BZY the lowest.

### Distinct local adaptation patterns across landforms

To identify genomic signatures of local adaptation in *F. danxiaensis*, we combined two complementary analytical approaches (RDA and LFMM) (Figure 5a,b; Figure S10-12) with high-resolution soil and climate data. Comparative analysis of 20 soil variables between DL and KL revealed significant differences in 14 variables, among which 13 had significantly higher concentrations in KL than in DL (Figure S7; Table S15). Among the 19 climatic variables compared, 14 showed significant differences with 10 factors significantly higher in the KL (Figure S8; Table S16). These results indicate that the environmental condition in DL is more harsh compared to KL, which could potentially impose greater selective pressures on *F. danxiaensis*. Totally, 18 environmental variables were selected for RDA based on pairwise correlation results, including total Potassium (TK), Available Potassium (AK), Available Phosphorus (OP), Total Phosphorus (TP), Total Magnesium (TMg), Available Nitrogen (AN), Total Calcium (TCa), Available Zinc (AZn), Available Iron (AFe), Bio17, Bio15, Bio12, Bio10, Bio9, Bio7, Bio5, Bio4, and Bio1) (Figure 5a; Figure S9). All results of RDA in three groups (all populations, landform group, and lineage group) showed statistically significant (p < 0.001) results with corrected R^2^ values. Environmental variables explained 39.12% of genetic variations across all populations of *F. danxiaensis* (Figure 5a). Within the landform group, environmental variables accounted for 31.96% and 17.19% of the genetic variation in the DL and KL populations (Figure S10a, b). A total of 12,577 and 16,500 candidate variants, involving 6,093 and 7,723 genes that potentially associated with local adaptation were identified in the DL and KL populations, respectively. Additionally, based on LFMM analysis (p < 1e–8), we found 16,778 variants, involving 7,664 genes in the DL populations, in contrast to only 3,057 variants, involving 1,690 genes in the KL populations (Figure S12a).

**Figure 4.**
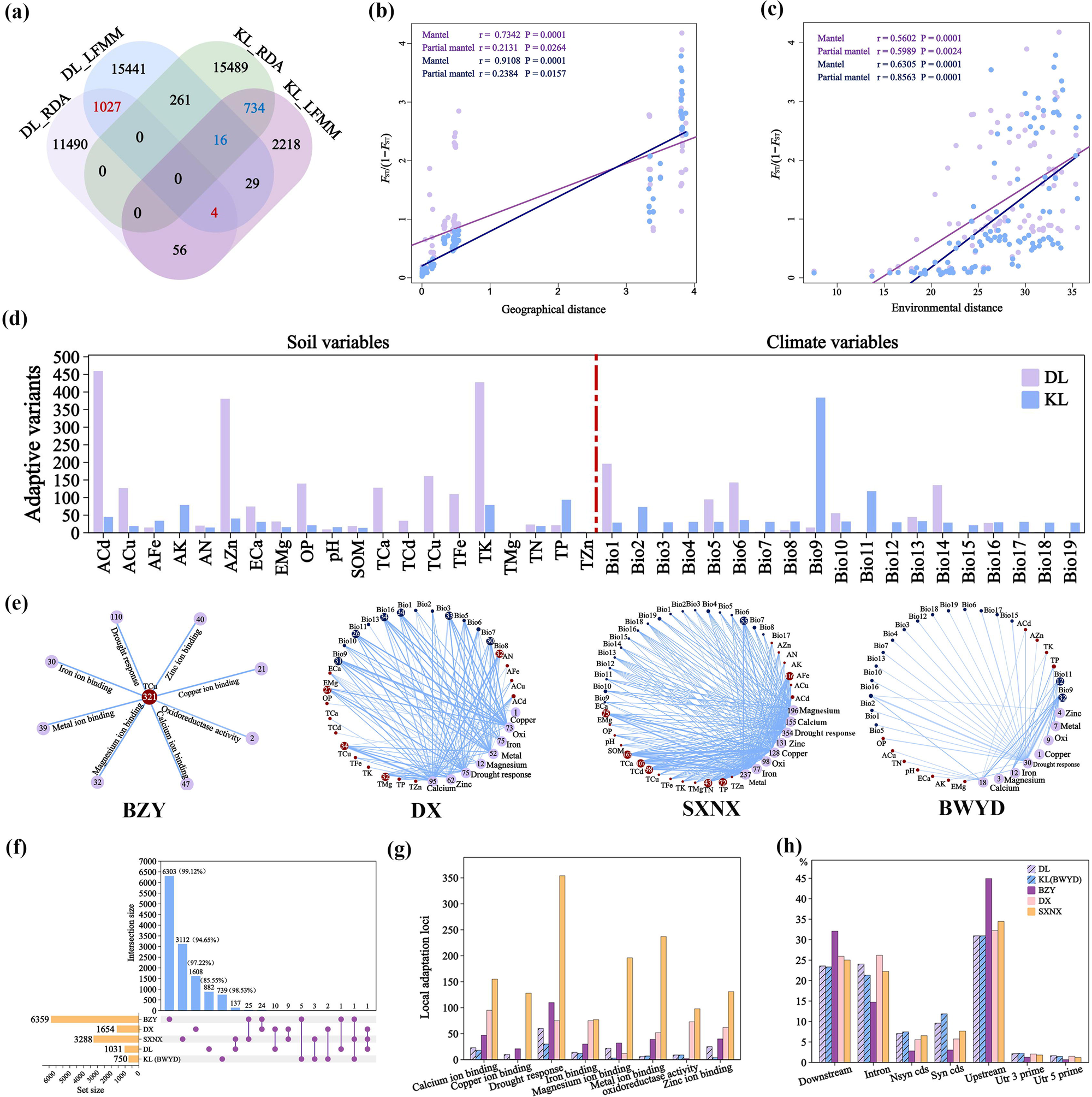
Patterns of local adaptation in *Firmiana danxiaensis* across landforms and genetic lineages. (a) Venn diagram of the Redundancy analysis (RDA) and Latent Factor Mixed Model (LFMM) analysis results for the Danxia landform (DL) and Karst landform (KL) populations. (b,c) Isolation by distance (IBD) and isolation by environment (IBE) analyses (Mantel test, two-sided) in DL populations based on neutral (blue dots/line) and local adaptive variants (purple dots/line). (d) Bar chart showing the influence of 39 environmental variables (20 soil variables and 19 climatic variables) on adaptive variants in DL and KL populations. (e) Environmental-genetic network across four genetic lineages. Numbers within blue and dark red circles represent variants significantly associated with environmental variables. Numbers within purple circles indicate variants categorized by function. (f) Upset plot showing overlap of core local adaptive variants across landforms and genetic lineages. (g) Bar plot of the number of adaptive variants associated with key Gene ontology functions. (h) Proportional distribution of adaptive variants across the genome.

**Figure 5.**
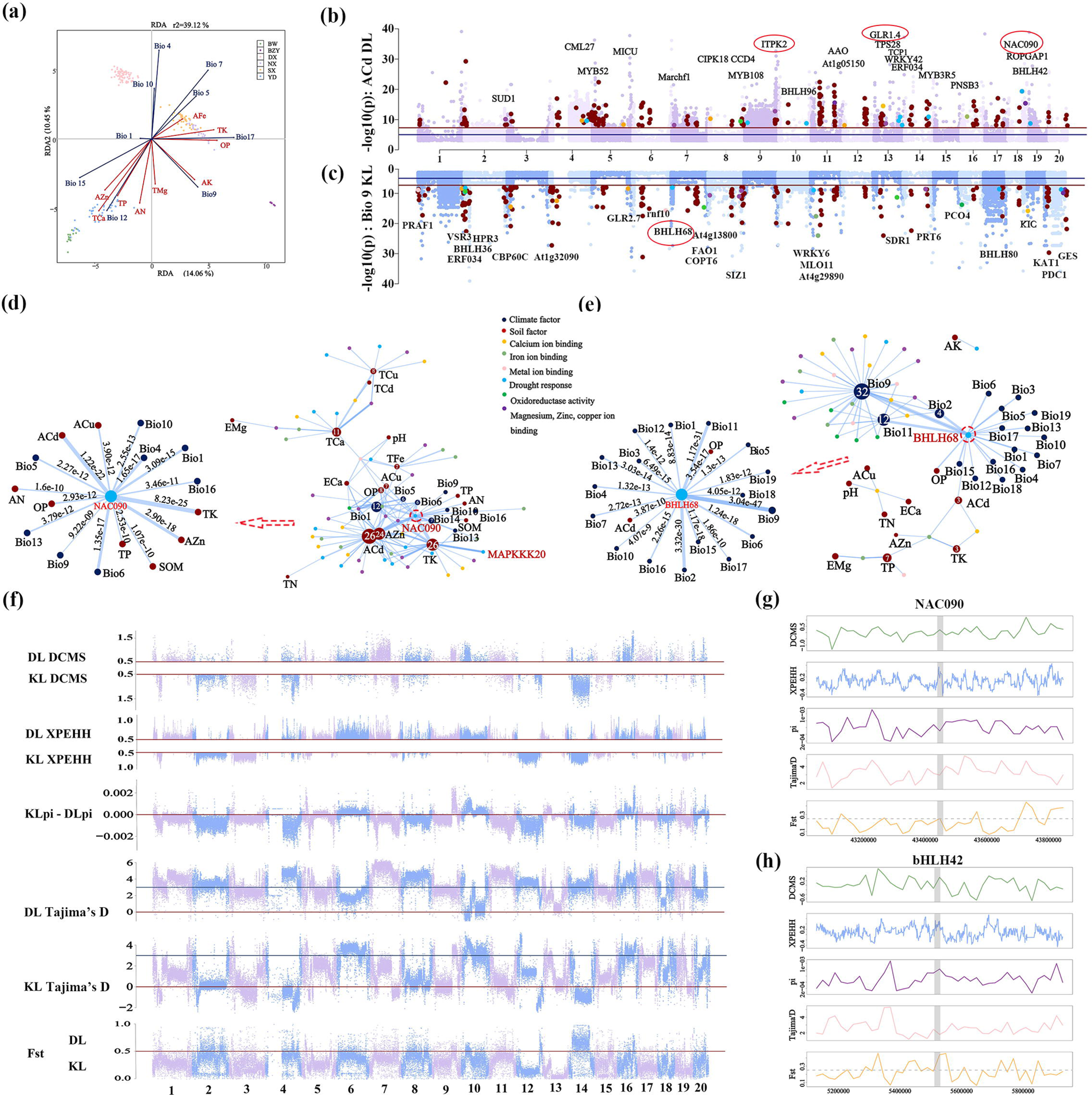
Signatures of local adaptation and positive selection in *Firmiana danxiaensis* across landoforms. (a) Redundancy analysis (RDA) of all populations of *F. danxiaensis*. Dots represent individuals, colored by sampling sites. Proximity among dots reflects similarity in genetic composition. Blue arrows indicate climatic variables; red arrows indicate edaphic variables. (b, c) Manhattan plots of variants associated with available cadmium (ACd) in the Danxia landform (DL) populations (b) and mean temperature of the driest quarter (Bio9) in the Karst landform (KL) populations. Horizontal lines represent significance thresholds: blue (p = 1e-5) and red (p = 1e-8). Red points indicate putative local adaptive variants. Annotated genes at significant variants are color-coded by function, consistent with d,e. The genes in the red circles represent those that are significantly influenced by environmental factors. (d, e) Network relationships between environmental variables and adaptive variants with key funcitons in the DL (d) and KL (e) populations. Edge thickness corresponds to the significance level (p-value); thicker lines indicate stronger associations. (f) Manhattan plots of three genetic statistic scores (pi, Tajima’s D, and Fst), Cross-Population Extended Haplotype Homozygosity (XP-EHH) values, and decorrelated composite of multiple signals (DCMS) scores between DL and KL populations. (g, h) A zoom-in on three genetic statistic scores (pi, Tajima’s D, and Fst), XP-EHH values, and DCMS scores, for the *NAC090* gene and *bHLH42* gene regions and their upstream and downstream 400 kb extension. Each genetic statistic is based on a sliding window analysis using nonoverlapping 20 kb windows.

To further pinpoint core variants associated with local adaptation, we integrated the results of RDA and LFMM analyses to identify overlapping variants. There were 1,031 and 750 core variants, involving 685 and 531 genes for local adaptation in DL and KL populations, respectively. Notably, no shared variants were found between the two landform populations (Figure 4a). This support that *F. danxiaensis* developed distinct adaptation patterns in two contrasting edaphic islands. Additionally, environment exerted a stronger influence than geography on adaptive variation in the DL populations, as indicated by Mantel and partial Mantel tests (Figure 4b, c). This was reflected in a clear pattern of IBE for both adaptive and neutral variants. In contrast, the pattern of IBD was weak after correcting for environmental effects, despite being significant in standard Mantel tests. For the KL populations, neither adaptive nor neutral variants exhibited significant IBD or IBE in either Mantel or partial Mantel tests (Figure S13).

Furthermore, to gain a deeper understanding of local adaptation patterns, we mapped core local adaptive variants to the LFMM analysis results. The results revealed a complex network between environmental variables and local adaptive variants (Figure 5d, e; Figure S14). In the DL populations, multilocus, multifactor interaction accounted for 91.30% of the adaptive variants (Table S17). Edaphic variables showed a more significant impact (74.91%) than climatic variables (Figure 4d; Table S17), with key environmental variables being Available Cadmium (ACd), TK, and AZn (Figure 4d, 5b, 5d; Figure S14a; Table S17). To identify the primary selective pressures, we performed Gene Ontology (GO) functional enrichment analysis on the core adaptive variants. Drought response was the most enriched GO terms for local adaptation, followed by metal ion binding functions related to Zn, Ca, Mg, and Fe ions (Figure 4g, 5d; Table S18, S19). Notably, *NAC090*, was significantly influenced by 16 environmental variables (8 climatic and 8 edaphic variables), with TK (p = 8.23e-25), ACd (p = 1.22e-22), and AZn (p = 2.90e-18) showing the strongest effects (Figure 5d). Additionally, *MAPKKK20*, contained seven adaptive variants, all significantly influenced by TK (Figure 5d). For the KL populations, the complex interaction accounted for 89.72% of adaptive variants (Figure 5e; Figure S14b; Table S20). In contrast, climatic variables exhibited a stronger (67.10%) influence than edaphic variables (Figure 4d, 5e; Table S20). The main variables were Bio9 (Mean temperature of driest quarter) and Bio11 (Mean temperature of coldest quarter) (Figure 4d, 5c, 5e; Figure S14b). Again, drought response was the most enriched GO term, influenced by 23 environmental variables, followed by Ca and Fe ion binding (Figure 4g, 5e; Table S21, S22; Note that the BWYD lineage and KL populations are the same). Among these genes, *BHLH68* was significantly influenced by 19 environmental variables (17 climatic and 2 edaphic variables), with Bio9 (p = 3.04e-47), Bio11 (p = 1.17e-31), and Bio2 (p = 3.32e-30) exhibiting the strongest associations (Figure 5e).

To further assess the selection pressures acting on local adaptive variants, we integrated five complementary metrics — XP-EHH, *F*_ST_, π, Tajima’s *D*, and DCMS statistic (Ma *et al*., 2015; Cao *et al*., 2023)(Figure 5f) — to better comprehensively identify genomic signals of local adaptation. For the XP-EHH results, variants with signatures of positive selection were enriched in Chr 6 and Chr 10 in the DL populations, whereas in the KL populations, positively selected variants were concentrated on Chr 2, 3, 12, and 14 (Figure 5f; Figure S15b). Consistently, the selective sweep results of *F*_ST_, π, Tajima’s *D*, and the DCMS statistic revealed a highly convergent chromosomal distribution of positive selection signals across these same genomic regions (Figure 5f). Furthermore, we integrated the results from both GEA and XP-EHH analyses by focusing on overlapping variants that exhibited signatures of both local adaptation and positive selection. There were 130 adaptive variants (12.61%) that showed signatures of positive selection (Figure S15d) in DL populations. Among these, six genes were functionally linked to key adaptive traits, including drought response (*MAPKKK20*, *NAC090*, and *BHLH42*), magnesium ion binding (*ITPK2*, *GES*), and calcium ion binding (*GLR1.4*) (Figure 5g, h; Figure S16), indicating their crucial roles in adaptation to DL. In contrast, only six local adaptive variants (0.8%) (Figure S15d) exhibited evidence of positive selection in the KL populations. These results support that local adaptation of *F. danxiaensis* is likely driven by the polygenic selection (Fagny & Austerlitz, 2021; Yeaman, 2022).

### Distinct local adaptation patterns across lineages

To estimate the local adaptation patterns among the four identified lineages of *F. danxiaensis*, we applied the same methods as those used at the landform level. Environmental variables explained a lower proportion of genetic variation (BZY: 1.97%. DX: 3.95%. SXNX: 15.28%. BWYD: 17.19%) (Figure S10c-e), and the number of potential local adaptive variants varied across lineages based on RDA and LFMM results (Figure S12b; Table S23). Core local adaptive variants were largely lineage-specific, with minimal sharing among the four lineages (Figure 4f), indicating long-term evolution shaped by geographic isolation and divergent demographic trajectories.

The adaptive variants of BZY were only linked to a single edaphic variable—total copper (TCu). In contrast, other three lineages exhibited complex network between adaptive variants and environmental variables, accounting for 96.21% and 98.37% of the adaptive associations in DX and SXNX lineages, respectively (Figure 4e; Figure S17). Once more, drought response was the most enriched GO term across all four lineages, followed by functions related to metal ion binding, although each lineage exhibited distinct functional profiles (Figure 4e, g).

In addition, the distribution of the core local adaptive variants exhibited similar characteristics across both landforms and lineages, with about 10% located in coding regions (Figure 4h; Table S24). In addition, we identified 1,977 (31.09%), 132 (7.98%), and 91 adaptive variants (2.77%) that displayed signatures of positive selection in BZY, DX, and SXNX lineages, respectively (Figure S15d). BZY exhibited the strongest signatures of positive selection, with the highest XP-EHH values and the largest number of positively selected variants, followed by SXNX, DX, and BWYD (Figure S15a,b, S18). Notably, five chromosomes (Chr 2, Chr 6, Chr 9, Chr 12, and Chr 13) exhibited either the highest or the lowest numbers of positively selected variants in 12 pairwise comparisons (Figure S15b, S18). Specially, Chr 2 showed the fewest selected variants, while Chr 6 had the most in SXNX lineage. In DX, Chr 9 displayed the most, and in BWYD, Chr 12 had the highest number. In BZY, all chromosomes except Chr 13 showed more signals of positive selection (Figure S15b, S18). Similarly to adaptive variants, most of positively selected variants were located in non-conding regions across both landform and lineages (Figure S15c).

## Discussion

### Adaptability characteristic of *F. danxiaensis* genome

Genome size, a fundamental attribute of species, is increasingly recognized as an important factor in adaptation to environmentally stressful habitats (Samatadze *et al*., 2023). This is exemplified by species such as *Helianthus annuus*, whose 3 Gb genome underlies a robust tolerance to drought, salt, and diverse soil types (Badouin *et al*., 2017), and *Opisthopappus taihangensis*, whose large genome is likely key to its specialization in the nutrient-poor, barren cliffs of the Taihang Mountains (Wang *et al*., 2025). In line with this, the adaptability of *F. danxiaensis* may also be distinguished by its relatively expansive genome size (1.51 Gb). Within the Malvaceae family, the genome of *F. danxiaensis* notably exceeds that of several sequenced relatives, including *D. zibethinus* (715 Mb) (Teh *et al*., 2017), *B. ceiba* (895 Mb) (Gao *et al*., 2018), *H. littoralis* (819 Mb) (Wang *et al*., 2024), *C. capsularis* (339 Mb) (Islam *et al*., 2017), and *T. cacao* (327 Mb) (Argout *et al*., 2011). Thus, *F. danxiaensis*, which inhabits the shallow, erosion-prone, and metal-rich soils of DL and KL, may utilize its large genome as a crucial adaptive strategy to thrive in these harsh edaphic island environments.

WGD is prevalent in plants and serves as a major evolutionary force promoting genomic diversification and adaptation (Soltis *et al*., 2014). Under environmental stress, WGD may enhance phenotypic plasticity through functional redundancy and neofunctionalization, enabling rapid adaptation to climatic fluctuations (Andersson & Hughes, 2009). Such genomic events have been shown to buffer against selective pressures and increase long-term survival, as exemplified in lineages like *Opisthopappus* during the Eocene–Oligocene cooling (Bielski *et al*., 2018; Wang *et al*., 2025). In line with these observations, *F. danxiaensis* has also undergone two WGD events. The earlier one, at ∼133 Ma, corresponds to the core eudicot-wide γ event, which contributed to gene family expansion and functional diversification (Jiao *et al*., 2011). A more recent WGD occurred around 20 Ma, coinciding with the early Miocene—a period marked by Asian monsoon formation and global cooling (Zachos *et al*., 2001). These events likely furnished *F. danxiaensis* with the genetic repertoire necessary to adapt to the metal-rich, drought-prone soils of DL and KL edaphic islands.

TEs, particularly LTR–RTs, serve as a major source of genomic innovation and have been shown to facilitate lineage-specific adaptation through mechanisms such as insertional mutagenesis and gene duplication (Warren *et al*., 2015; Oliver *et al*., 2013; Niu *et al*., 2019). These elements are known to respond to various external stimuli, including abiotic stresses such as freezing, thereby contributing to genome reorganization under environmental challenges (Galindo-González *et al*., 2017; Huang *et al*., 2021). Among Malvaceae species, *F. danxiaensis* exhibits one of the highest documented proportions of LTR–RTs (67%), surpassed only by *F. yunnanensis* (75%), compared to ∼26% in durian and ∼50% in *B. ceiba* and *Heritiera* species (Teh *et al*., 2017; Gao *et al*., 2018; Wang *et al*., 2024; Yang *et al*., 2024). Notably, approximately 40% of LTR–RTs insertions in *F. danxiaensis* occurred between 1∼2.5 Ma (Figure 1c), a period that coincides with the Quaternary glacial–interglacial cycles. These repeated climatic oscillations likely imposed strong selective pressures, necessitating rapid evolutionary adaptations via genomic restructuring—a process in which TE activation and LTR insertion may have played a critical role.

In addition, TFs involved in stress responses are often preferentially retained following WGD events, with these duplicated genes contributing not only to developmental regulation but also to adaptation under abiotic stress (Wu *et al*., 2020; Khan *et al*., 2018). Among the key TF families known to be associated with drought response— such as *C2H2*, *ERF*, *MYB*, *NAC*, *WRKY*, *bHLH*, and *bZIP* (Shinozaki & Yamaguchi-Shinozaki, 2007; Hao *et al*., 2021)—several show pronounced expansion in *F. danxiaensis* compared to general species (Figure 1f). This pattern of lineage-specific TF expansion mirrors observations in other edaphic specialists such as *Primulina huaijiensis* (Feng *et al*., 2020b), supporting the view that these TFs may have contributed to the adaptation of *F. danxiaensis* to the challenging environments of DL and KL. In summary, the evolution of *F. danxiaensis* has been characterized by two WGD events, substantial accumulation of LTR–RTs, and expansion of seven drought responses-related TF families. These genomic signatures have collectively shaped its relatively large genome and likely underpin its genomic basis of adaptation .

### Interplay of genetic drift and divergent selection

On edaphic islands, population differentiation is often driven by the joint effects of genetic drift and divergent selection. Genetic drift tends to be strong in spatially isolated populations due to founder effects, small effective population size (Ne), demographic bottlenecks, and limited gene flow (Franks, 2010; Frankham, 1998; Eldridge *et al*., 1999). In contrast, divergent selection becomes prominent where environmental heterogeneity exists among islands (Weigelt *et al*., 2013), as observed in various edaphic specialists (Cao *et al*., 2023; Feng *et al*., 2020; Arnold *et al*., 2016). The balance between these forces is modulated by demographic and genetic connectivity: lower Ne and restricted gene flow enhance drift and weaken selection (Charlesworth, 2009), whereas higher gene flow and larger population sizes mitigate drift and help maintain adaptive variation (Charlesworth, 2009; Charlesworth & Willis, 2009; Keller & Waller, 2002).

Our study of *F. danxiaensis* provides empirical support for this theoretical framework. The species exhibited pronounced genetic structure, with the easternmost Danxia lineage (BZY) showing the strongest signals of genetic drift—including the lowest genetic diversity, highest inbreeding, and lowest gene flow—consistent with long-term isolation and small Ne. In contrast, the BWYD, DX, and SXNX lineages, which experienced less geographical isolation and higher gene flow, retained greater genetic diversity and were less affected by drift, despite historical bottlenecks (Fig. 3a). These results underscore how variation in connectivity and demography among edaphic islands can alter the interplay of stochastic and selective forces, ultimately shaping divergent evolutionary pathways within a single species.

### Multi-Level genomic signature of local adaptation to contrasting edaphic islands

Local adaptation results from the complex interplay between selective pressures and demography, particularly in spatially structured landscapes. In small, isolated populations, genetic drift can overwhelm selection, limiting the fixation of adaptive alleles and constraining evolutionary pathways (Leal *et al*., 2021; Ostridge *et al*., 2025; Wright, 1931). Conversely, populations with larger effective sizes and higher gene flow retain greater standing genetic variation, thereby enhancing adaptive potential (Bridle & Vines, 2007). In *F. danxiaensis*, we observed lineage- and landform-specific patterns of local adaptation, contrasting with the species-level signals often reported in the literature (Lazic *et al*., 2024; Marková *et al*., 2023; Sang *et al*., 2022). This lineage-level structuring reflects how divergent environmental conditions impose distinct selective pressures, leading to differentiated adaptive trajectories (Savolainen et al. 2007; Zhang et al. 2020). The more stressful environment in DL (Figure S7, S8) likely imposed stronger selection, resulting in more adaptive variants (1,031) than in KL (750) (Figure 4a). Furthermore, demographic differences reinforced these patterns: the BZY lineage—characterized by the smallest effective population size and lowest gene flow—exhibited a simplified adaptive architecture consistent with drift-dominated evolution. In contrast, southern lineages (BWYD, DX, SXNX) maintained higher levels of standing variation and showed clearer signatures of selection. These findings underscore the need for multi-level analyses—spanning landforms and genetic lineages—to fully resolve the genomic basis of local adaptation in edaphic specialists.

Furthermore, integrating genomic and environmental data has significantly advanced the identification of locally adaptive loci, a crucial step for deciphering the spatial distribution of adaptive variation and informing evidence-based conservation (De Mita *et al*., 2013; Armstrong *et al*., 2021; Ostridge *et al*., 2025). Yet, most previous studies have relied predominantly on selective sweep scans or GEA methods (Lazic *et al*., 2024; Sang *et al*., 2022; Wang *et al*., 2017; Arnold *et al*., 2016; Cao *et al*., 2023). For example, research on serpentine or karst adaptation in edaphic specialists such as *Arabidopsis arenosa* and *Platycarya longipes* used selective sweep approaches to detect localized adaptive variants without incorporating soil or climate data (Cao *et al*., 2023a; Arnold *et al*., 2016). In another representative case, GEA analysis of European beech (*Fagus sylvatica*), based solely on climatic variables, recovered a set of candidate loci—including one high-confidence region associated with winter temperature (Lazic *et al*., 2024). Such examples reflect a recurring methodological focus on either selection signatures or climate-driven adaptation, often omitting the integration of fine-scale edaphic variation. By contrast, our study among the first to combine high-resolution field-measured soil profiles with climatic variables to dissect the mechanisms of local adaptation in edaphic specialists. This integrated approach revealed a fundamental divergence in adaptive drivers: soil properties acted as the dominant selective force in DL populations, whereas climate exerted stronger influence in KL systems (Figure 4d; 5d, e). These results underscore the essential and often underappreciated role of spatially heterogeneous soil conditions in shaping adaptation within edaphic islands. Importantly, our field-based soil metrics enabled the detection of genomic signatures of adaptation that would have remained undetected in climate-only models.

At the molecular level, drought stress represents a major environmental challenge for plants in terrestrial ecosystems, particularly in edaphic islands where water availability is often limited (Fan *et al*., 2021; Huang *et al*., 2016). Plants have evolved sophisticated molecular mechanisms to cope with water deficit conditions, with TFs playing central roles in regulating drought response pathways (Thirumalaikumar *et al*., 2018; Jin *et al*., 2014). Among key regulators, *NAC* TFs have been extensively documented as crucial mediators of drought adaptation. Studies across multiple species demonstrate that *NAC* genes enhance drought tolerance through various mechanisms: *OsSNAC1* from rice improves drought resistance in cotton by promoting root growth and reducing transpiration (Liu *et al*., 2014), *OsNAC066* in rice reduces water loss and ROS accumulation (Yuan *et al*., 2019), while *MuNAC4* from *Macrotyloma uniflorum* enhances drought tolerance through improved root development and antioxidative defense (Pandurangaiah *et al*., 2014). Similarly, *bHLH* TFs contribute significantly to drought adaptation, often functioning within the ABA signaling pathway—a central regulator of plant drought responses (Liu *et al*., 2015; Kim & Kim, 2006; McCourt. 1999). In *Arabidopsis*, *bHLH112* acts as a transcriptional activator that enhances drought tolerance (Yuan *et al*., 2008), while *bHLH129* and *bHLH92* are integral components of ABA-mediated stress signaling. Beyond TFs, *MAPK* cascades serve as critical signaling modules in drought perception and response (Wang *et at*., 2015). *MAPKKK*s, as pivotal components of these cascades, regulate drought stress through various mechanisms (Cardinale *et al*., 2002. Colcombet & Hirt, 2008. Jonak *et al*., 2002): in rice, DSM1 (a Group B Raf-like *MAPKKK*) enhances drought resistance via ROS scavenging, while GhRaf19 in cotton acts as a negative regulator (Jia *et al*., 2016; Ning *et al*., 2010). Additional *MAPKKKs* including *OsMAPKKK28* and *OsMAPKKK8* have been associated with drought tolerance (Rao *et al*., 2010). In edaphic specialists, genomic analyses consistently reveal enrichment of drought-responsive functions (Arnold *et al*., 2016; Cao *et al*., 2023b; Feng *et al*., 2020b; Tao *et al*., 2016). Studies in karst-adapted species have reported expansion of *WRKY* and *bZIP* gene families known to enhance drought and salinity tolerance (Rushton *et al*., 2010; Singh *et al*., 2002; Wu *et al*., 2009), as observed in *Primulina huaijiensis* (Feng *et al*., 2020). Our findings in *F. danxiaensis* align with these patterns, showing significant signals of local adaptation and positive selection in *NAC*, *bHLH*, *WRKY*, *bZIP*, *TCP*, and *DREB* TF families (Table S11), which are established regulators of drought response (Hao *et al*., 2021; Shinozaki & Yamaguchi-Shinozaki, 2007). Notably, key genes with the strongest environmental associations—including *NAC090*, *MAPKKK20*, and *bHLH68* (Figure 5d, e)—all may play established roles in drought adaptation (Shinozaki & Yamaguchi-Shinozaki 2007; Sun *et al*., 2017; Hao *et al*. 2021). The significant signatures of local adaptation and positive selection observed at *NAC090*, *bHLH42*, and *MAPKKK20* in DL populations highlight their putative importance in adaptation to the drought-prone Danxia landform.

In addition to drought stress, metal ion homeostasis represents another major selective pressure in edaphic island systems (Cao *et al*., 2023; Tao *et al*., 2016; Arnold *et al*., 2016). Previous studies on karst adaptation have highlighted calcium tolerance as a key factor, identifying *TPC1* as an important candidate gene (Cao *et al*., 2023; Tao *et al*., 2016). Although *TPC1* has also been associated with serpentine adaptation in *Arabidopsis arenosa* (Arnold *et al*., 2016), we did not detect significant adaptive signals at this locus in *F. danxiaensis*. This discrepancy may reflect methodological differences—such as the use of outlier tests in previous studies versus GEA in ours—or distinct evolutionary paths in different edaphic systems. Instead, our GEA results indicate that local adaptation in *F. danxiaensis* involves a coordinated response to multiple metals, including Ca, Fe, Cu, Zn, and Mg (Figure 4e, g; 5d, e), indicating a broader mechanism of ion homeostasis beyond calcium-specific tolerance. This multi-metal adaptive strategy is further supported by the significant enrichment of metal ion binding functions among expanded gene families (Figure S3).

In summary, this study demonstrates that parallel local adaptation in *F. danxiaensis* is driven by the combined effects of drought stress and multi-metal ion imbalances, mediated through lineage- and landform-specific genetic architectures. By integrating high-resolution environmental data with genomic analyses across multiple organizational levels—from landforms to genetic lineages — we have uncovered how the balance between selection and drift shapes adaptive divergence in edaphic islands. The distinct selective regimes between Danxia and Karst landforms, coupled with the identification of key candidate genes such as *NAC090*, *bHLH42*, and *MAPKKK20*, provide a mechanistic understanding of adaptation to extreme soil conditions. Future functional studies on the identified candidate genes will be crucial to fully elucidate the molecular mechanisms underlying adaptation in these unique and threatened ecosystems.

### Conservation implications

The genomic patterns identified in this study provide direct guidance for conserving *F. danxiaensis*. The northern BZY lineage, characterized by critically low nucleotide diversity (π = 4.94 × 10⁻⁵), high inbreeding (mean *F*_ROH_ = 0.50), and elevated genetic load, should be designated as a high-priority conservation unit. To counter inbreeding depression and boost evolutionary potential, assisted gene flow from the SXNX lineage—which shares the Danxia landform and exhibits the highest genetic diversity (π = 8.21 × 10⁻⁴)—is recommended as a genetic rescue measure (Tallmon *et al*., 2004; Frankham, 2015; Whiteley *et al*., 2015). Conservation strategies should further reflect landform-specific adaptation mechanisms (Ostridge *et al*., 2025). In Danxia sites, efforts should focus on mitigating heavy metal stress (e.g., ACd, TK, AZn) and restoring soils, whereas Karst conservation should emphasize climate resilience, particularly thermal adaptation (Bio9, Bio11), supported by long-term monitoring and genetic-offset analyses. Broader interventions—including habitat restoration, pollution control, and ecological monitoring—should address the principal selective pressures of drought and multi-metal ion stress common to both landforms. Collectively, these findings affirm the need for genomics-based, lineage-tailored approaches to preserve the evolutionary potential of edaphic specialists under environmental change.

## Materials and Methods

### Genome assembly and annotation

Seeds were collected from the natural range of *F. danxiaensis* (114°12′22″ E, 25°07′14″ N) in Nanxiong city, Guangdong Province, China for karyotype analysis to determine the chromosome number. A total of 75 Gb of subreads (50 × depth) from the PacBio Sequel platform and 76 Gb of short reads (50 × depth) from the Illumina HiSeq platform were generated for *F. danxiaensis*. The genome was initially *de novo* assembled and then polished by four rounds of Illumina short reads. The genome size was estimated through the analysis of 150 bp paired-end reads, computation of 17 bp *K*-mer frequencies using Jellyfish v2.3 (Marçais & Kingsford, 2011). Additionally, flow cytometry analysis was performed for further validation. The karyotype analysis revealed that *F. danxiaensis* have a chromosome base of 20 (2*n* = 40). The improved contigs were further assembled into scaffolds with a scaffold N50 of approximately 69 Mb. Hi-C library was prepared and anchored to 20 chromosomes following the standard procedure (Zhang *et al*., 2019). A total of 150 Gb of Hi-C data (100 × depth) were generated using the Illumina platform. BUSCO (Simão *et al*., 2015) with a plant database of 248 conserved plant genes was used to estimate the completeness of the assembly. We annotated protein-coding genes and gene structures by integrating *de novo* identification, homology-based prediction, and RNA-Seq data. This information was then combined into a nonredundant gene model set using EVidenceModeler (EVM v1.1.1) (Haas *et al*., 2008). For function annotation, we annotated the protein-coding genes against the SwissProt (Schneider *et al*., 2004), KEGG (Kanehisa & Goto, 2000), Nr (http://www.ncbi.nlm.nih.gov/protein), Pfam (Mistry & Finn, 2007), InterPro (https://www.ebi.ac.uk/interpro/), and GO (Gene Ontology) (Ashburner *et al*., 2000) databases. We also identified a set of non-coding RNAs.

### Comparative genomics analyses

To identify key genomic features associated with the adaptation of *F. danxiaensis*, we performed a series of comparative genomic analyses. Repetitive sequences were identified using a combination of homology-based prediction and de novo methods. Tandem repeats were detected using the *ab initio* approach with TRF v4.09 (http://tandem.bu.edu/trf/trf.html). We identified LTRs with the LTR_retriever method. Specifically, LTR_finder (Xu & Wang, 2007) and LTRharvest (Ellinghaus *et al*., 2008) were used to identify all the existing LTR sequences in the *F. danxiaensis* genome. Following clustering and filtering, we estimated insertion times (T) on the basis of T = K/2r, where K is the genetic distance and r is the substitution rate (1.3×10^−8^).

The protein sequences of 15 plant species were used to cluster gene families, including seven from the Malvaceae family, *F. danxiaensis*, *Heritiera littoralis* (Wang *et al*., 2024), *Durio zibethinus* (Teh *et al*., 2017), *Gossypium arboreum* (Wang *et al*., 2022), *Bombax ceiba* (Gao *et al*., 2018), *Theobroma cacao* (Argout *et al*., 2011), *Corchorus capsularis* (Zhang *et al*., 2021), and eight outgroup plants, *Arabidopsis thaliana* (The Arabidopsis Genome Initiative, 2000), *Platycarya longipes* (Cao *et al*., 2023), *Solanum lycopersicum* (Tomato Genome Consortium, 2012), *Marsdenia tenacissima* (Zhou *et al*., 2023), *Primulina huaijiensis* (Feng *et al*., 2020b), *Zea mays* (Chen *et al*., 2022), *Oryza sativa* (Kawahara *et al*., 2013), and *Amborella trichopoda* (AMBORELLA GENOME PROJECT *et al*., 2013). We used OrthoFinder (Emms & Kelly, 2019) to cluster gene families and identified 331 single-copy gene families across 15 species for further analysis. The phylogenetic tree for 15 species was constructed using the maximum likelihood (ML) method in RAxML v8.2.12 (Stamatakis, 2014), and species divergence times were estimated with the MCMCTREE program in PAML (Yang, 2007). Ten calibration nodes were applied to calibrated the divergence times (http://www.timetree.org). We then used CAFÉ v5 (Mendes *et al*., 2021) to analyze the expansion and contraction of gene families. Using conditional likelihood as the test statistic, we calculated the p values for each lineage, with a p value of 0.01 considered significant. Kyoto Encyclopedia of Genes and Genomes (KEGG) and Gene ontology (GO) enrichment analyses were performed on the expanded and contracted gene families to determine their functions.

Protein sequence alignments were conducted for seven species (*F. danxiaensis*, *D. zibethinus*, *G. arboreum*, *H. littoralis*, *P. huaijiensis*, *C. capsularis*, and *M. tenacissima*). We appiled MCscanX (Wang *et al*., 2012), followed by multiple sequence alignment with Muscle to calculate 4dTv values. Ks values were then estimated using the codeml mode of PAML v4.9(Yang, 2007).

Transcription factors (TFs) in the *F. danxiaensis* genome were identified using iTAK (Zheng *et al*., 2016), and classified based on the PlnTFDB (Pérez-Rodríguez *et al*., 2010) and PlantTFDB (Jin *et al*., 2014) databases, with categories having a frequency <10 removed. The ranking of each TF category was calculated, and the Kolmogorov-Smirnov test in SPSS was applied to assess normal distribution. The data were then standardized, and z-scores for each category were computed (z-score = (x–μ)/ɕ). Besides, TFs from 36 eudicots were calculated and compared with those of *F. danxiaensis* (Feng *et al*., 2020b) (Table S13). Finally, functional annotation of the over expressed TFs in *F. danxiaensis* was performed.

### Plant sample collection, sequencing, and variant calling

A total of 225 individuals from 20 populations (15 Danxia landform (DL) populations and five Karst landform (KL) populations) of *F. danxiaensis* were sampled for the whole-genome resequencing. Paired-end libraries with a 150 bp insert were sequenced on Illumina HiSeq instruments by NovoGene (Beijing, China). We mapped all reads to *F. danxiaensis* reference genome with default settings implemented in BWA-mem2 v2.2 (Li & Durbin, 2010). The variation detection was performed using GATK v4.3.0 (McKenna *et al*., 2010) with HaplotypeCaller algorithm and default filter parameters. Further quality control was applied based on missing rate (> 5%), minor allele frequency (MAF< 0.05), Linkage Disequilibrium (r^2^ > 0.2), and Hardy Weinberg Equilibrium (P < 1e-5).

### Soil sample collection, processing, and measurement

To obtain high-resolution soil data, we collected soil samples from sites of all 20 *F. danxiaensis* populations. At each site, five random points, each more than five meters apart, were selected to collect soil samples using soil core sampler (0-25 cm depth). After natural air-drying, debris such as dead branches, leaves, crop roots, and stones were removed. The soil samples were first ground with a mortar and pestle, then sieved through a 0.149 mm mesh. A total of 20 variables were measured using standard procedures as below. Soil pH was determined using a glass electrode with a water-to-soil ratio of 2.5:1. Soil organic matter (SOM) was quantified by external heating potassium dichromate oxidation. Total nitrogen (TN) was measured via the Kjeldahl digestion-distillation titration method. Total phosphorus (TP) and total potassium (TK) were determined using sodium hydroxide fusion followed by molybdenum antimony colorimetry and flame atomic absorption spectrophotometry, respectively. Available nitrogen (AN) was assessed by alkaline hydrolysis diffusion. Available phosphorus (OP) was extracted using hydrochloric acid-ammonium fluoride and measured by molybdenum antimony colorimetry, while available potassium (AK), calcium (ECa), magnesium (EMg), and cadmium (ACd) were extracted with ammonium acetate and analyzed by flame atomic absorption spectrophotometry. Total calcium (TCa), magnesium (TMg), and iron (TFe) concentrations were determined by flame atomic absorption spectrophotometry. Available iron (AFe) was measured using the o-phenanthroline colorimetric method. Total cadmium (TCd) was quantified by graphite furnace atomic absorption spectrophotometry. Total copper (TCu) and zinc (TZn), as well as available zinc (AZn) and copper (ACu), were measured by inductively coupled plasma optical emission spectrometry (ICP-OES), with available metals extracted by diethylene triamine pentaacetic acid (DTPA).

### Population structure and population genetic parameters

Population genetic structure was inferred using Admixture v. 1.3.0 (Alexander *et al*., 2009), with *K* values set from 2 to 10. The *K* value where the CV curve began to smooth was selected. Principal component analysis (PCA) was performed using Plink v 1.9 (Purcell *et al*., 2007). In addition, a phylogenetic tree of 225 samples was constructed using IQ-TREE v 2.2.0.3 (Minh *et al*., 2020) with 1,000 replicates for bootstrap confidence analysis. Nucleotide diversity (π), Tajima’s D and the fixation index (*F*_ST_) were calculated for the high-quality SNPs in 20 kb nonoverlapping windows using vcftools v0.1.16 (Danecek *et al*., 2011).

To compare the difference in inbreeding among *F. danxiaensis* lineages, we applied vcftools v0.1.16 (Danecek *et al*., 2011) to calculate heterozygosity (Het) for individual of *F. danxiaensis*. Inbreeding coefficients (*F*_IS_) were then derived based on individual heterozygosity. The Plink v 1.9 (Purcell *et al*., 2007) software was used to identify runs of homozygosity (ROH) longer than 500 kb. The proportion of ROH in genome (FROH) was calculated. Besides, linkage disequilibrium (LD) values were calculated using PopLDdecay v3.42 (Zhang et al. 2019). The R package fasteprr (Gao *et al*., 2016) was employed to calculate the recombination rate (r) across 50 kb sliding windows.

### Demography history and gene flow

To infer historical changes in the effective population size (Ne) of *F. danxiaensis*, we applied Fitcoal v1.1 (Hu *et al*., 2023) based on site frequency spectrum (SFS). We used non-CDS SNP sites for analysis to mitigate the effects of purifying selection (Hu *et al*., 2023). The Python script easySFS.py (Gutenkunst *et al*., 2009) was then employed to calculate the SFS. We defined 10 years as one generation for *F. danxiaensis*, set the mutation rate at 7.5×10^-6^ kb/generation, repeat the analysis 100 times, and applied a –logLPRate of 1.3%. Additionally, referred to the differentiation history of *F. danxiaensis* obtained from Fitcoal, we set up the simulation scenario for Fastsimcoal2 (Excoffier *et al*., 2013). Each scenario was repeated 10 times, and finally, the analysis was run 100 times based on the optimal parameters. The mutation rate was set the same as in the Fitcoal analysis. The AIC value was then computed using the formula AIC = 2k–2ln (ML), where k is the number of parameters and ML is the maximum likelihood. A lower absolute AIC value indicates a better model fit, with the model being closer to the actual one.

To determine gene flow among *F. danxiaensis* lineages, allele frequencies for each cluster were calculated by Plink v 1.9 (Purcell *et al*., 2007). Treemix enables the analysis of gene flow direction. We applied Treemix v1.12 (Pickrell & Pritchard, 2012) and constructed a maximum likelihood tree with 100 SNPs per unit, varying m from 1 to 10, repeated 5 times with 1000 bootstraps, and setting k to 100 while calculating the standard deviation. Subsequently, the plot_optM function from the R package OptM (Fitak, 2021) was employed to identify the optimal number of migration events using the Evanno method. Finally, the migration patterns were visualized by built-in script plotting_funcs.R. Meanwhile, we applied Dsuite v0.5 r50 (Malinsky *et al*., 2021) to quantify the gene flow among *F. danxiaensis* populations, identifing P1, P2, P3, and outgroup populations based on the maximum likelihood tree results without migration constructed by Treemix. The D-statistics (Patterson *et al*., 2012), p-values, Z-values, and f4 ratios were calculated using the Dtrios module, with p-values adjusted using the R package ’qvalue’ (Storey, 2003). The fb values were then calculated using the Fbranch command (Malinsky *et al*., 2018), which assigns gene flow evidence to specific internal branches of the phylogenetic tree, with visualization performed by the built-in script dtools.py. Furthermore, the results from the Dtrios module were filtered for combinations of P1, P2, and P3 with p < 1e–4 and Z > 3. These combinations were then analyzed in the Dinvestigate module to calculate fd statistics (Martin *et al*., 2015) using built-in ’ABBABABAwindows.py’ script for non-overlapping genomic windows of three sizes (10 kb, 50 kb, and 100 kb), which quantify the proportion of the genome affected by gene flow (Fu *et al*., 2022).

### Genotype**-**Environment Association (GEA) analyses

To reduce false positives in GEA analyses, we applied two different approaches to identify environment-associated variants across landforms and lineages of *F. danxiaensis*. Environmental data include 20 edaphic variables (Table S15) and 19 climatic variables (Fick & Hijmans, 2017) (Table S16) from 20 populations. First, we used a redundancy analysis (RDA)(Capblancq & Forester, 2021) to identify genetic variants showing a significant strong relationship with multivariate environmental axes. Variables used in study were selected based on pairwise correlations analysis (|r| < 0.65). The RDA includes three groups: the overall populations (225 samples), the landform group (DL group (171 samples), and the KL group (54 samples), and the lineage group (BZY (23 samples), DX (76 samples), SXNX (72) samples), BWYD (54 samples (same as KL))). The R^2^ values in the analysis were adjusted using the RsquareAdj command, and the significance of the results was assessed through 999 permutations via the anova command. The explained variance for the first two axes was then calculated based on the adjusted R^2^. To identify SNPs significantly influenced by environmental variables, the average and standard deviation of all SNPs across the first three axes were computed. SNPs with values exceeding |mean ± 3SD| were classified as potential local adaptative variants (Forester *et al*., 2016).

Second, we applied a univariate latent factor linear mixed model (LFMM) implemented in the R package lfmm (Frichot *et al*., 2013) to identify associations between allele frequencies and the 39 environmental variables. Based on the number of ancestry lineages inferred with Admixture, we conducted LFMM with four latent factors to account for population structure in the genotype data. The analysis encompassed three groups, which were consistent with those in the RDA. Following p-value correction, variants with p-values smaller than 1e-8 were selected as candidate local adaptive variants. The results for each environmental variable were exported, followed by counting and ranking the number of variants significantly linked to each variable. We then assessed the effect of these variables on genetic variation in DL and KL populations of *F. danxiaensis*. Based on the results from LFMM and RDA, the overlapping variants were identified as the core local adaptive variants. We then analyzed the types and proportions of these variants, and performed their localization and gene ontology analysis. Furthermore, to gain deeper insight into the local adaptation patterns, we aligned the core variants with the LFMM analysis results. The relationship between environmental variables and local adaptive variants was examined, including the number of variants significantly influenced by each environmental variable, as well as the number of environmental variables affecting each variant.

In addition, to explore and compare the influence of geography and environment on the genetic variation of adaptive and neutral variants, we employed Mantel and partial Mantel tests at landform level. These tests were used to examine the associations between (*F*_ST_/1–*F*_ST_) and geographic as well as environmental distances, with significance determined through 999 permutations using the R package vegan (Oksanen *et al*., 2025). Geographic distances were represented by Euclidean distance, while environmental distances were represented by Canberra distance.

To further assess the selective pressures acting on local adaptive variants, we performed a selective sweep analysis based on genome-wide SNP data. First, cross-population extended haplotype homozygosity (XP-EHH) was estimated on a per-SNP basis using Selscan 2.0 (Szpiech, 2021), following the approach of Sabeti *et al*. (2007). We then calculated Tajima’s *D*, nucleotide diversity (π), and the fixation index (*F*_ST_) in 20 kb sliding windows using VCFtools. To integrate these multiple signals and enhance the detection of selective sweeps, we combined XP-EHH, *F*_ST_, π, and Tajima’s *D* into a decorrelated composite of multiple signals (DCMS) statistic, computed in non-overlapping 20 kb windows as implemented in Ma *et al*. (2015) and Cao *et al*. (2023). Given that XP-EHH was estimated per SNP while the other metrics were window-based, we defined candidate positively selected regions using the top 5% of XP-EHH values for subsequent analysis to ensure consistency in signal resolution. Finally, candidate adaptive variants under positive selection were identified based on the overlap between GEA and XP-EHH signals.

## AUTHOR CONTRIBUTIONS

YLL, HFC, and JW conceived and designed the research. YLL, JCL and HFC collected plant materials. YLL performed data analysis, plotted manuscript Figs and drafted the manuscript. JCL, LD, RCZ, HFC, and JW revised the manuscript. All authors discussed the results and commented on the manuscript.

## ACKNOWLEDGEMENTS

We thank Prof. Song Ge (Institute of Botany, Chinese Academy of Sciences), Prof. Suhua Shi (Sun Yat-sen University), Prof. Ming Kang (South China Botanical Garden, Chinese Academy of Sciences), and Prof. Baosheng Wang (South China Botanical Garden, Chinese Academy of Sciences) for constructive comments, and Prof. Qiang Fan (Sun Yat-sen University) and Prof. Wenbo Liao (Sun Yat-sen University) for providing leaf material for YDD populations. We acknowledge Director Fang Chen and Director Zaixiong Chen of the Danxia Mountain Management Committee, Engineer Pingsheng Zhong of the Nanxiong Forestry Bureau, Section Chief Daowan Liu of the Shixing Forestry Bureau, Senior Engineer Yuanqiu Li of the Shimentai National Nature Reserve, Guangdong, Senior Engineer Xinyu Jia of the Guangdong Qingxin Baiwan Provincial Nature Reserve Management Office, Senior Engineer Minshui Lin of the Yong’ an Forestry Bureau, Fujian Province, Quan Zhong from the Taoyuandong Scenic Area in Yong’an City, Xianyuan Xu of the Dehua County Forestry Bureau, Dehua County, for their assistance during the field investigations and specimen collection. We thank Zhongcai Fan, Wentao Wang, Ting Wang, Mingxia Li, Yinling Chen, and Xiaohang Zhang for assistance with plant sample collection in Guangdong and Fujian Province, Zhongcai Fan, Mingchen Lin, Shuangwen Deng, Yinling Chen, and Xiaohang Zhang for soil sample collection, Dr. Chao Feng for assistance in genome assembly, Xuejing Jiang, Dongyi Chen, and Yong Tan for treatment of soil samples, and Yilan Chen for assistance with chromosome karyotype analysis. This work was supported by Guangdong Flagship Project of Basic and Applied Basic Research (grant: 2023B0303050001) and Self-deployed project of the South China Botanical Garden, Chinese Academy of Sciences (grant: E561150011).

## CONFLICT OF INTEREST STATMENT

The authors have no conflicts of interests to declare.

## DATA AVAILABILITY STATEMENT

The DNA sequencing data for the *Firmiana danxiaensis* genome assembly have been deposited in the NCBI database under BioProject PRJNA1261868 via the link https://dataview.ncbi.nlm.nih.gov/object/PRJNA1261868. The raw sequence data of 225 newly sequenced accessions have been deposited in the NCBI database under BioProject PRJNA1264699 via the link https://dataview.ncbi.nlm.nih.gov/object/PRJNA1264699. The data on environmental variables are from WorldClim (https://www.worldclim.org/).

## SUPPORTING INFORMATION

**Figure S1.** *K*–mer frequency distributions of *Firmiana danxiaensis* genome.

**Figure S2.** Flow cytometry histogram.

**Figure S3.** Gene ontology enrichment analysis of significantly expanded gene families in genome of *Firmiana danxiaensis*.

**Figure S4.** Cross-validation (CV) error values from ADMIXTURE analysis for *K* = 2 to 10

**Figure S5.** Genetic parameters among *Firmiana danxiaensis* lineages.

**Figure S6.** Heatmap of ABBA-BABA analysis.

**Figure S7.** Box plot comparing edaphic variables between Danxia and Karst landorms.

**Figure S8.** Box plot comparing climatic variables between Danxia and Karst landorms.

**Figure S9.** Correlation heatmap among 39 environmental variables.

**Figure S10.** Redundancy analysis of *Firmiana danxiaensis* across landforms and genetic lineages.

**Figure S11.** Manhattan plots of Latent Factor Mixed Model (LFMM) analysis results in populations of *Firmiana danxiaensis*.

**Figure S12.** Bar plot of LFMM results across landforms and genetic lineages of *Firmiana danxiaensis*.

**Figure S13.** Isolation by distance and isolation by environment analyses (Mantel test, two-sided) of *Firmiana danxiaensis* in Karst landform populations.

**Figure S14.** Network relationships between environmental variables and local adaptive variants across landforms of *F. danxiaensis*.

**Figure S15.** Signatures of positive selection of *Firmiana danxiaensis* across landforms and lineages.

**Figure S16.** Multinumerical line graphs of four key adaptive genes in Danxia landform populations.

**Figure S17.** Network relationships between environmental variables and local adaptive variants across lineages of *Firmiana danxiaensis*.

**Figure S18.** Upset plot of positively selected variants across landforms and lineages of *Firmiana danxiaensis*.

**Table S1.** Flow cytometry stats.

**Table S2.** Statistics of genome assembly for *Firmiana danxiaensis*.

**Table S3.** BUSCO evaluation results of *Firmiana danxiaensis* genome completeness.

**Table S4.** Structural annotation of inferred protein-coding genes in *Firmiana danxiaensis*.

**Table S5.** Functional annotation of predicted genes in *Firmiana danxiaensis*.

**Table S6.** Summary of repeat sequences identified in the *Firmiana danxiaensis* genome.

**Table S7.** Statistics of non-coding RNAs in the *Firmiana danxiaensis* genome.

**Table S8.** De novo identification and filtering results of LTR retrotransposons in the genome.

**Table S9.** Transcription factor gene families identified in *Firmiana danxiaensis*.

**Table S10.** Statistics of transcription factors across 36 plant species.

**Table S11.** Key transcription factors in Danxia and Karst populations of *Firmiana danxiaensis*.

**Table S12.** Summary statistics of Illumina resequencing data per individual.

**Table S13.** Fixation index among *Firmiana danxiaensis* populations.

**Table S14.** Likelihood comparison of demographic models for divergence history.

**Table S15.** Soil factor variables used in this study.

**Table S16.** Climatic environmental variables used in this study from WorldClim.

**Table S17.** Network between environmental variables and adaptive genes in Danxia landform populations.

**Table S18.** Core local adaptive genes in Danxia landform populations.

**Table S19.** Network between environmental variables and adaptive genes of key funciton in Danxia landform populations.

**Table S20.** Network between environmental variables and adaptive genes of key funciton in Karst landform populations.

**Table S21.** Core local adaptive genes in Karst landform populations.

**Table S22** Network between environmental variables and adaptive genes of key funciton in Karst landform populations.

**Table S23.** Statistics of local adaptive variants across landforms and genetic lineages.

**Table S24.** Genomic distribution of local adaptive variants in Danxia and Karst populations.

## Notes

### Competing Interest Statement

The authors have declared no competing interest.

