## supplementary figures for "Integrative genomic insights into parallel local adaptation to Danxia and Karst edaphic islands in the endangered tree *Firmiana danxiaensis*"

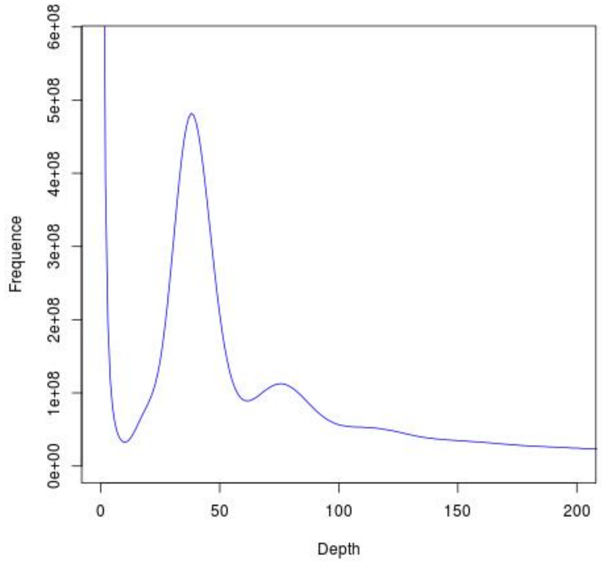


**Figure S1.** *K*–mer frequency distributions of *Firmiana danxiaensis* genome.

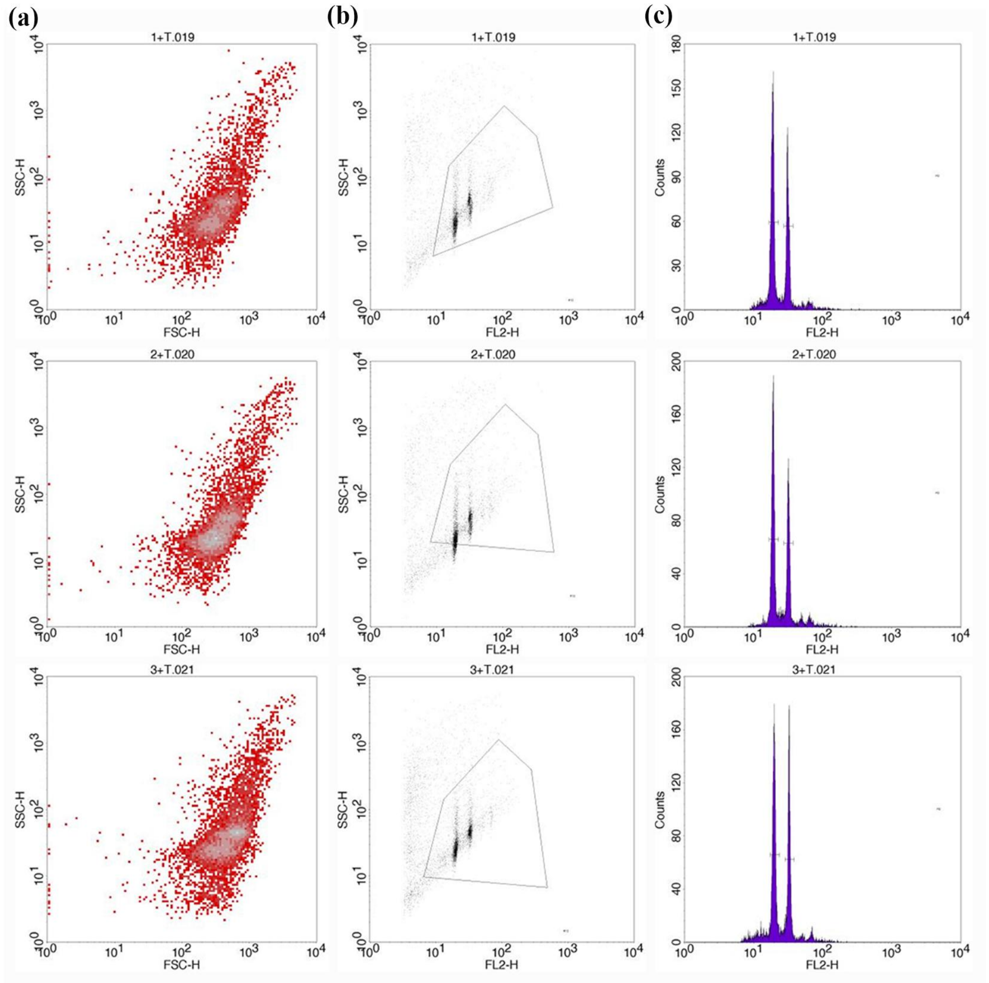


**Figure S2**. Flow cytometry histogram.

(a) Density plot of sample cell groups. (b) Fluorescence intensity dot plots. (c) Histograms.

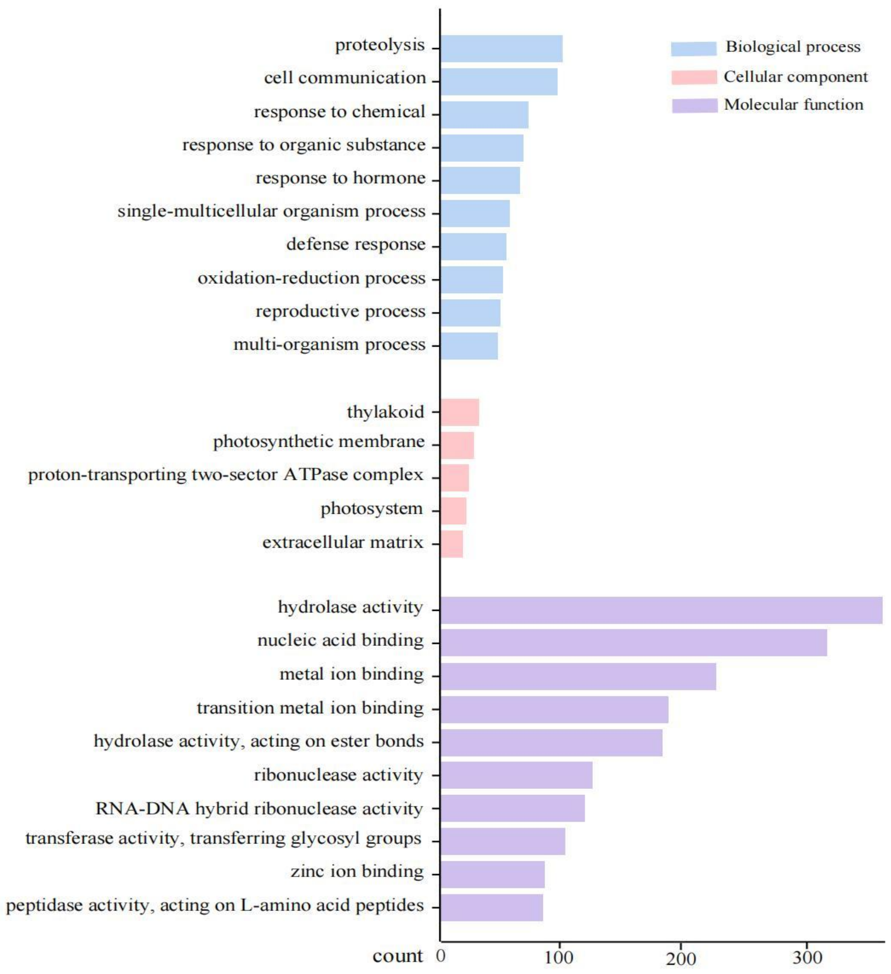


**Figure S3.** Gene ontology enrichment analysis of significantly expanded gene families in genome of *Firmiana danxiaensis*.

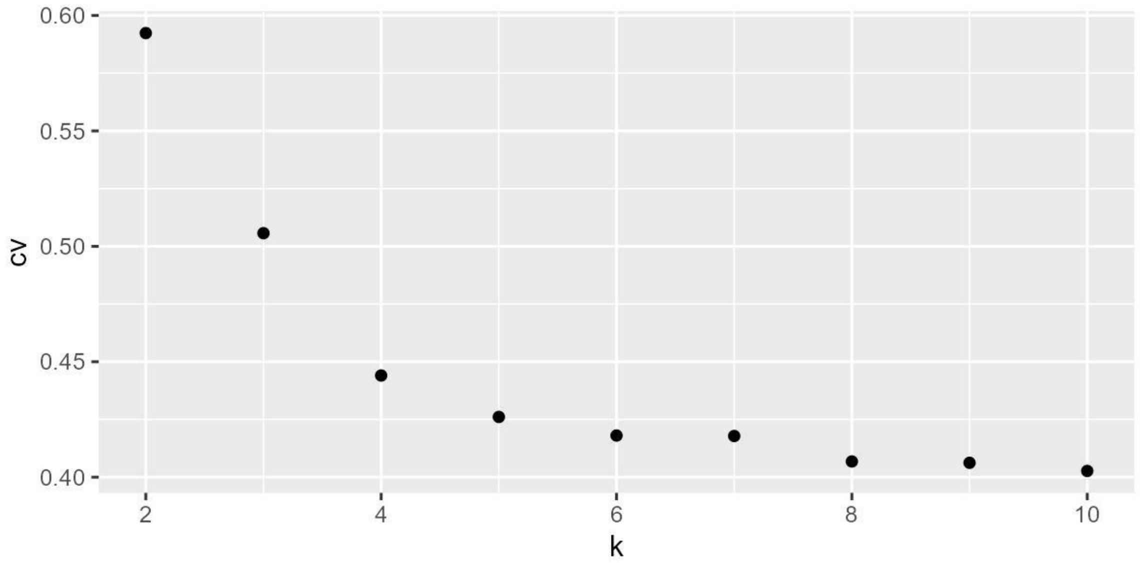


**Figure S4.** Cross-validation (CV) error values from ADMIXTURE analysis for $K = 2$ to 10.

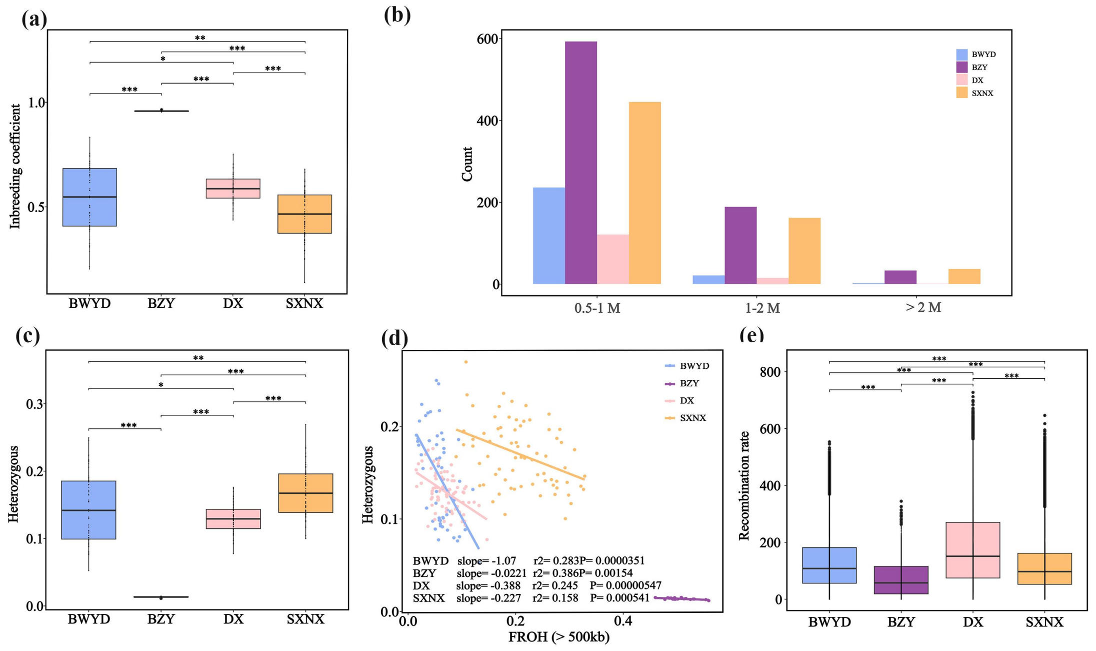


**Figure S5** Genetic parameters among *Firmiana danxiaensis* lineages.

(a) Inbreeding coefficient. (b) Bar chart of runs of homozygosity (ROH) across length groups. (c) Genome-wide heterozygosity. (d) Regression of FROH against heterozygosity. (e) recombination rates. (a, c, e) Center line = median. dots = outliers. *, P < 0.05. **, P < 0.01. ***, P < 0.001.

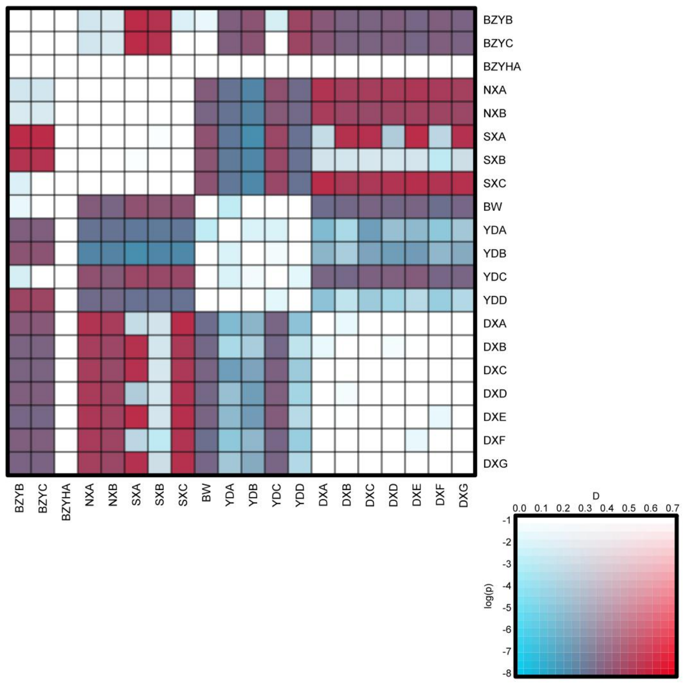


**Figure S6.** Heatmap of ABBA-BABA analysis.

Color represents D-value magnitude (warmer = larger); transparency reflects significance (less transparent = smaller p-values).

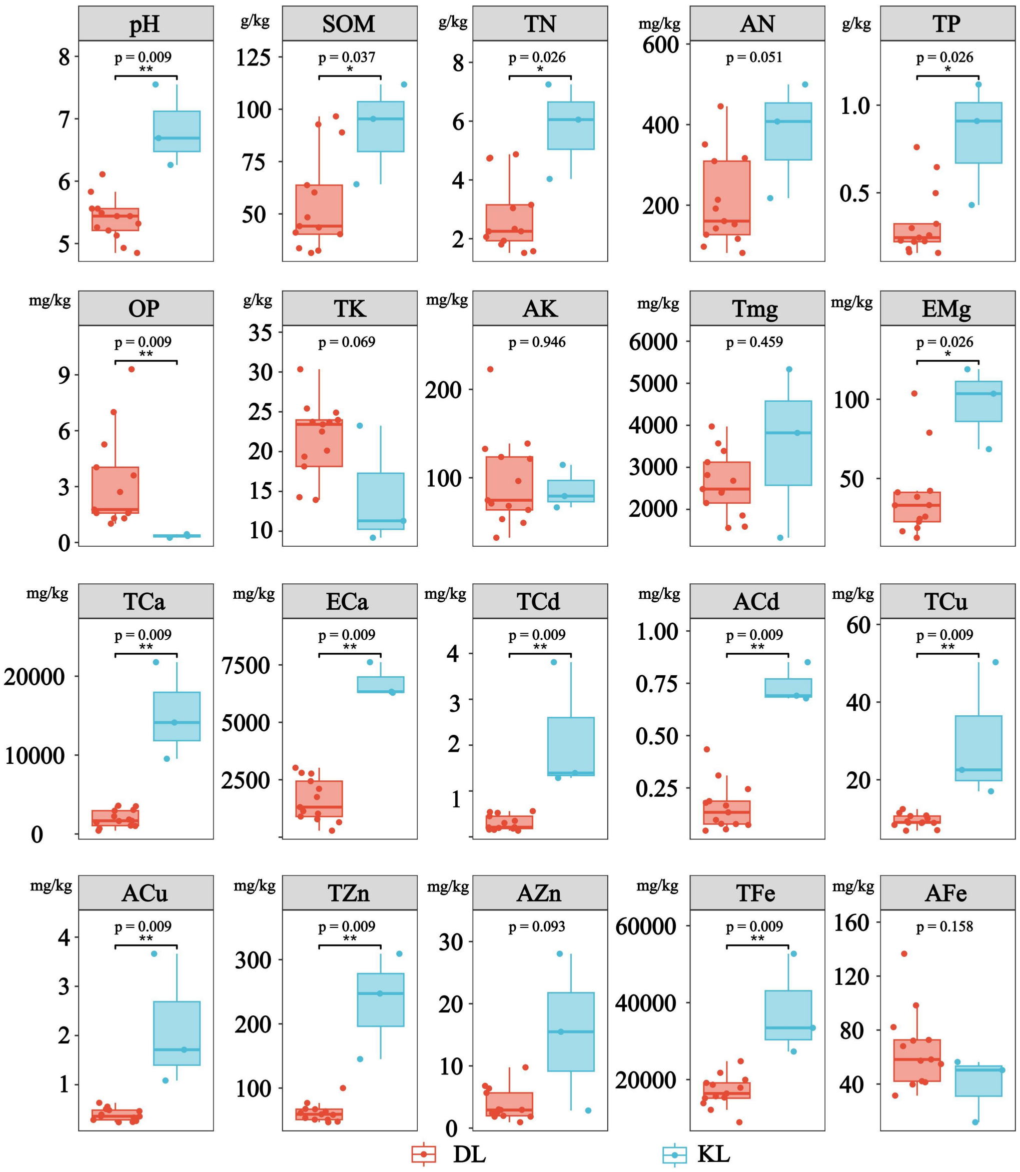


**Figure S7.** Box plot comparing edaphic variables between Danxia and Karst landorms. DL represents Danxia landform. KL represents Karst landform. *, P < 0.05. **, P < 0.01.

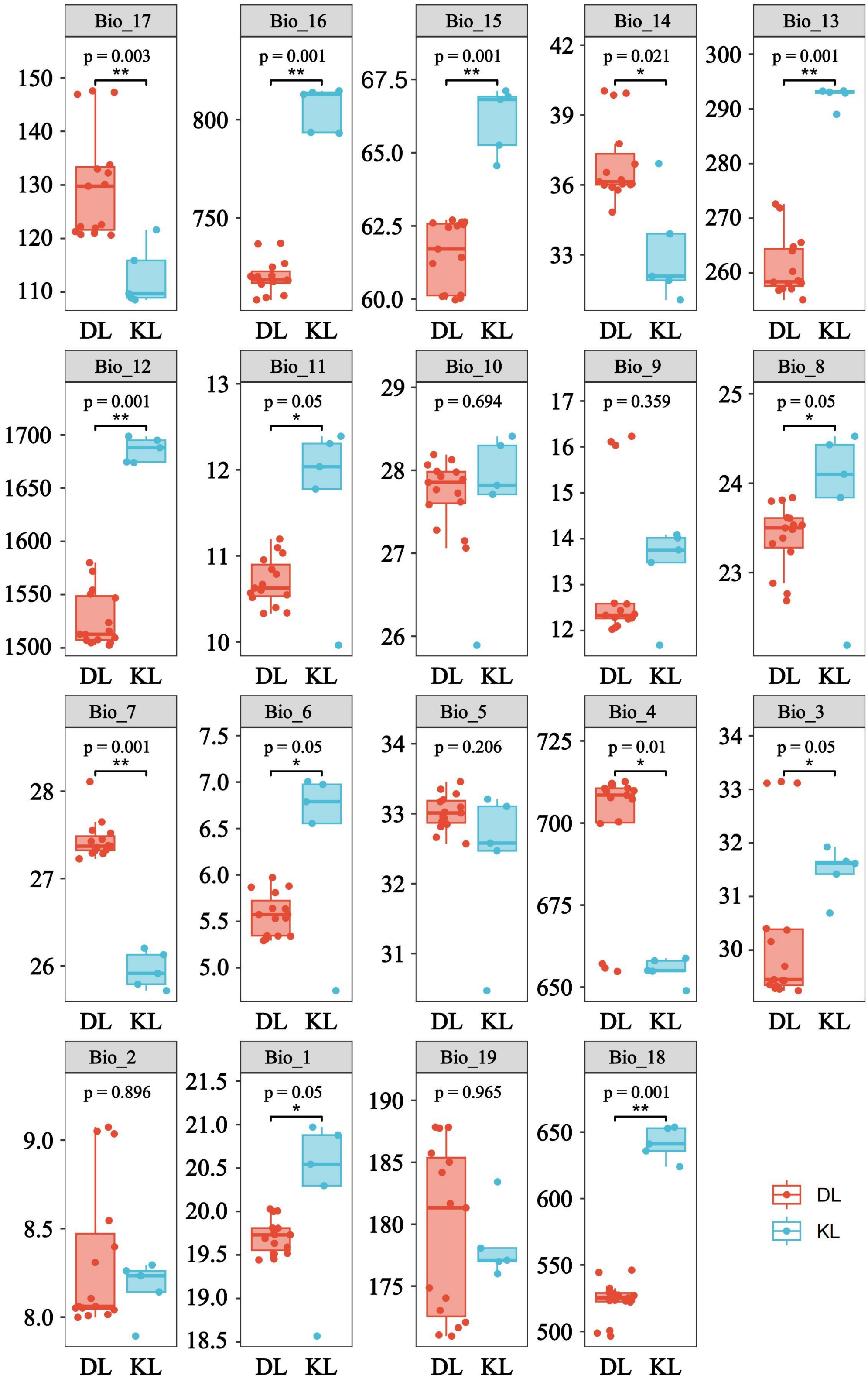


**Figure S8.** Box plot comparing climatic variables between Danxia and Karst landorms**.** DL represents Danxia landform. KL represents Karst landform. *, $P < 0.05$. **, $P < 0.01$.

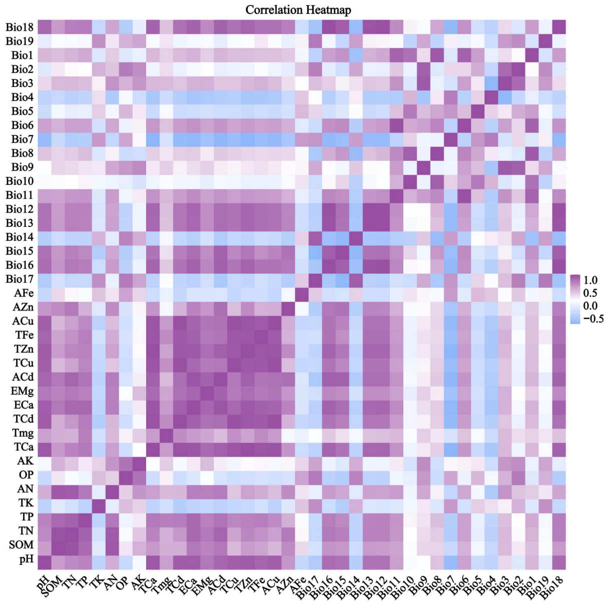


**Figure S9.** Correlation heatmap among 39 environmental variables.

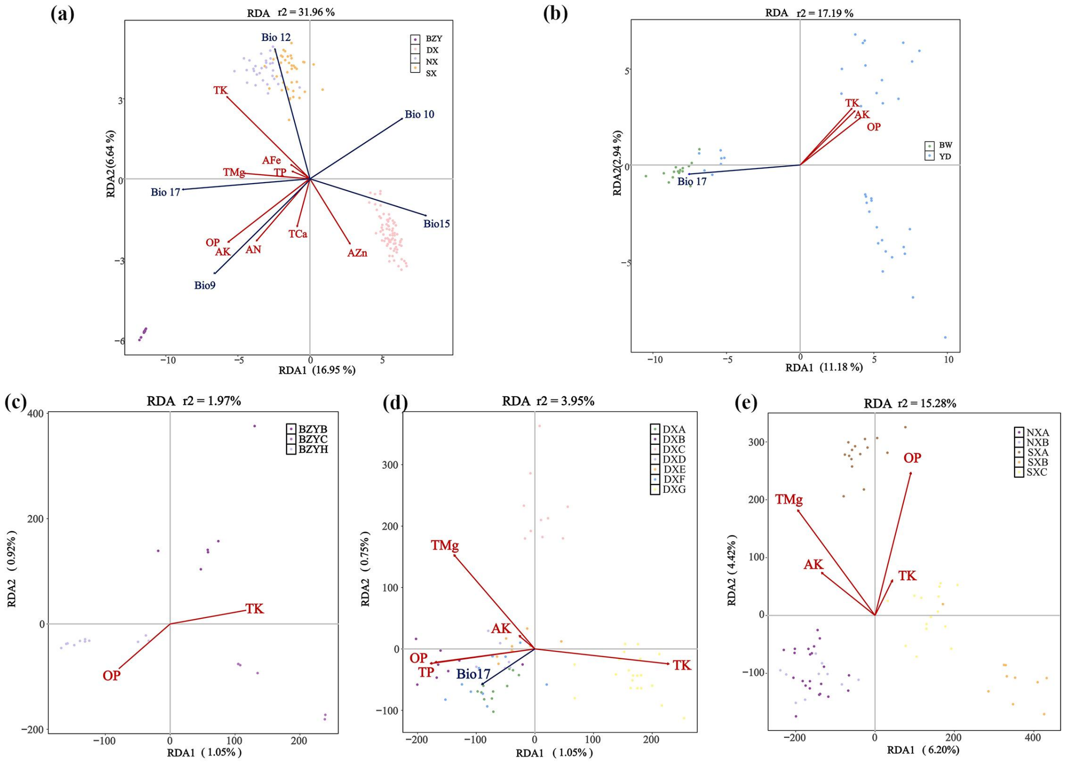


**Figure S10.** Redundancy analysis of *Firmiana danxiaensis* across landforms and lineages.

(a) RDA of the DL populations. (b) RDA of the KL (BWYD) populations. (c-e) RDA of the BZY (c), DX (d), and SXNX (e) lineages. Points represent individuals; different colors denote distributions. Blue arrows: climatic variables; red arrows: soil variables. Acute angle between arrows indicates positive correlation; right angle indicates no correlation; obtuse angle indicates negative correlation. Longer arrows represent stronger environmental influence. Arrow angles relative to axes indicate correlations with components; projection points reflect variable magnitude within populations. Axis labels indicate explained variance (%).

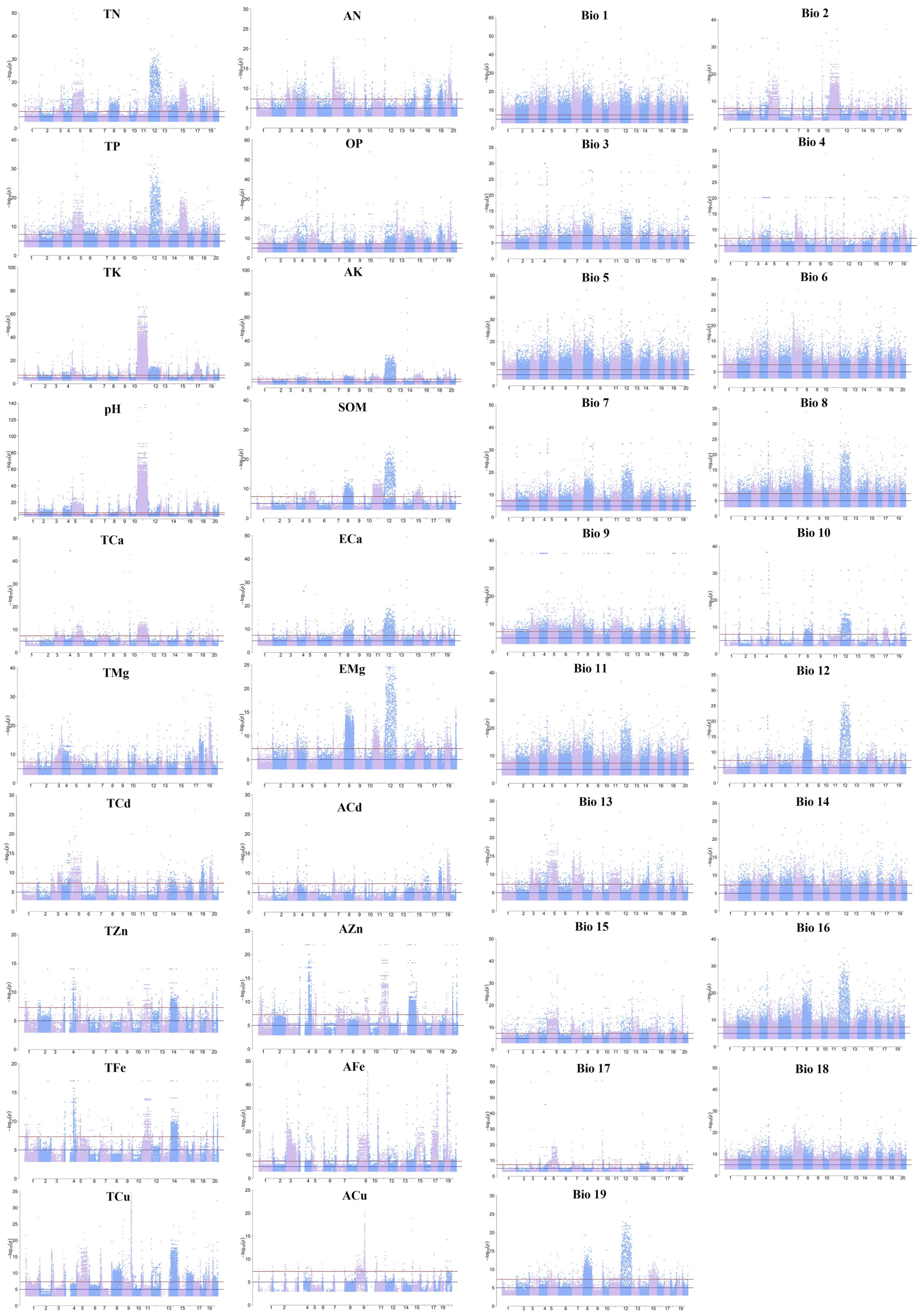


**Figure S11**. Manhattan plots of Latent Factor Mixed Model (LFMM) analysis results in populations of *Firmiana danxiaensis*.

Horizontal lines represent significance thresholds: blue (p = 1e-5) and red (p = 1e-8).

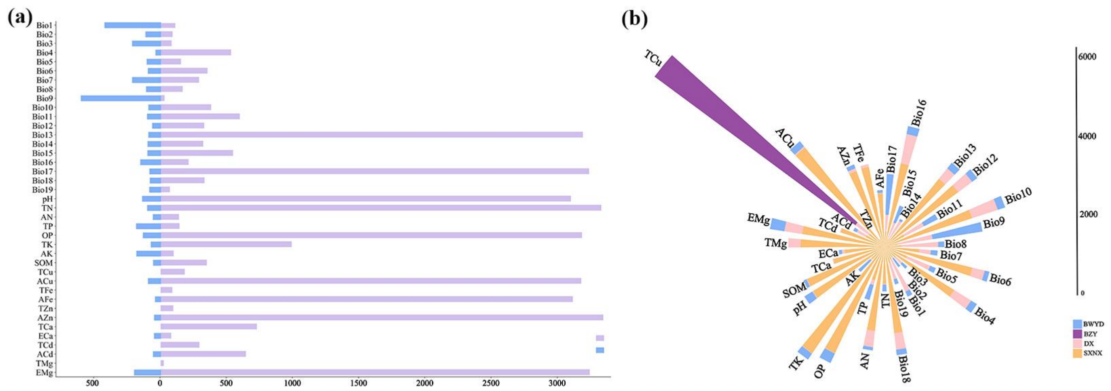


**Figure S12.** Bar plot of LFMM results across landforms and lineages of *Firmiana danxiaensis.*

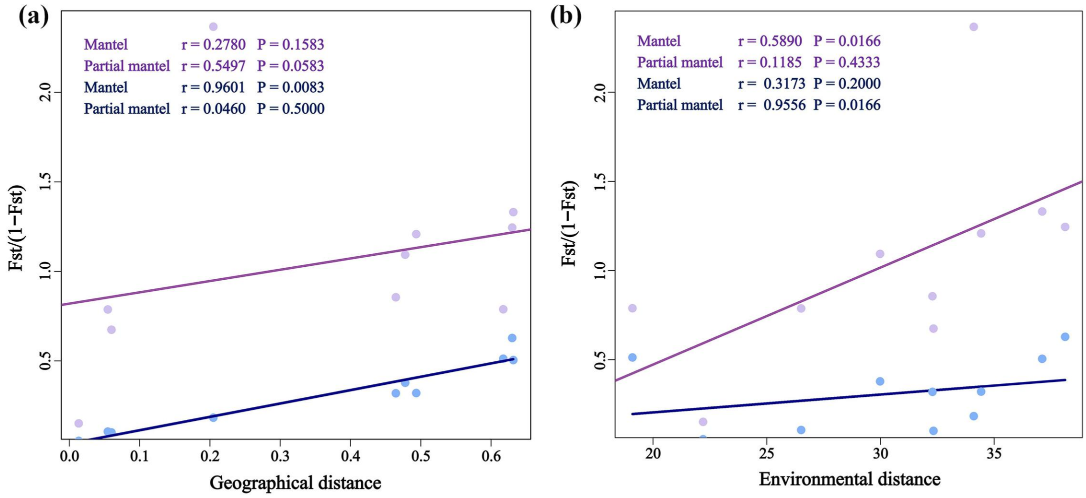


**Figure S13.** Isolation by distance and isolation by environment analyses (Mantel test, two-sided) of *Firmiana danxiaensis* in Karst landform populations.

(a-b) IBD and IBE analyses for KL populations (n = 54) based on neutral (blue dots/line) and local adaptation variants (purple dots/line).

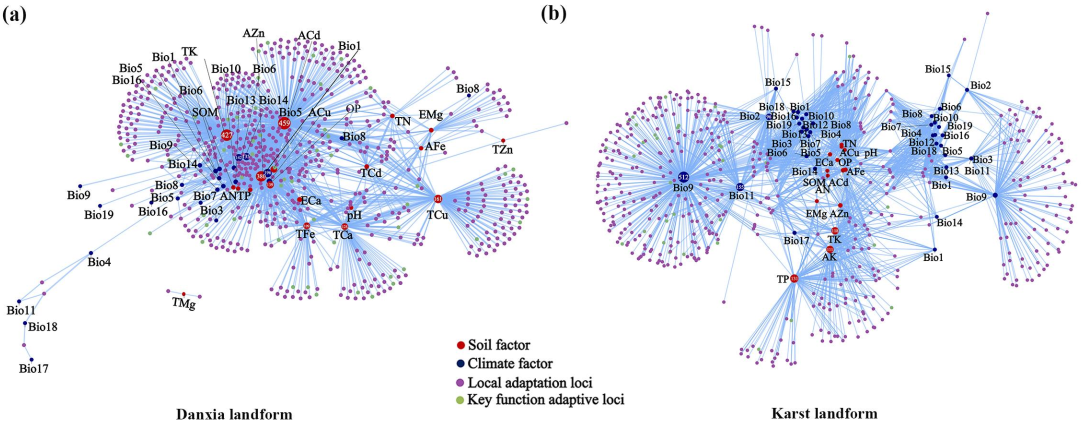


**Figure S14.** Network relationships between environmental variables and local adaptive variants across landforms of *Firmiana danxiaensis.*

Numbers in circles indicate variants significantly influenced by each environmental variable.

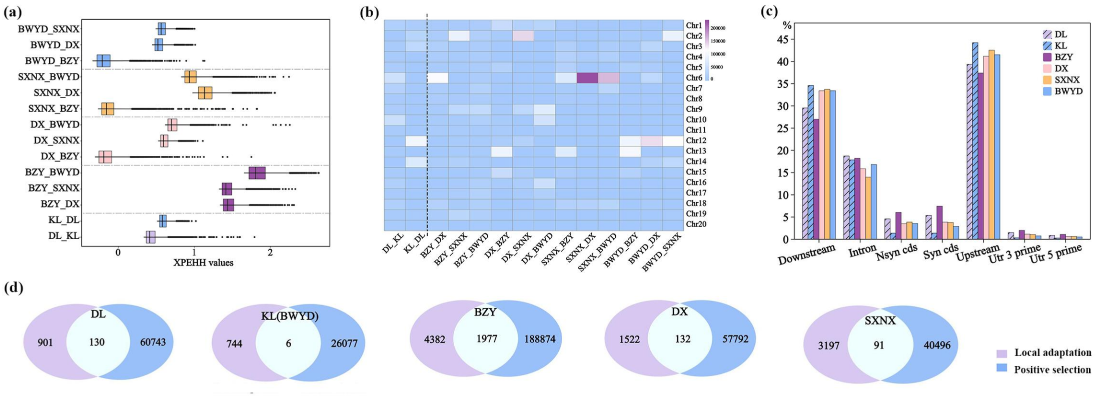


**Figure S15.** Signatures of positive selection of *Firmiana danxiaensis* across landforms and lineages.

(a) Box plot of cross-population extended haplotype homozygosity (XP-EHH) values in different pairwise comparisons. (b) Heat map showing the distribution of positively selected variants across 20 chromosomes for each pairwise comparison. (c) Proportional distribution of positively selected signatures across the genome. (d) Venn diagrams showing the overlap between signatures of local adaptation and positive selection. (a-d) BZY, DX, SXNX, and BWYD represent four lineages of *F. danxiaensis*. DL represents Danxia landform populations (BZY, DX, and SXNX). KL represents Karst landform populations, BWYD lineage is same as KL populations.

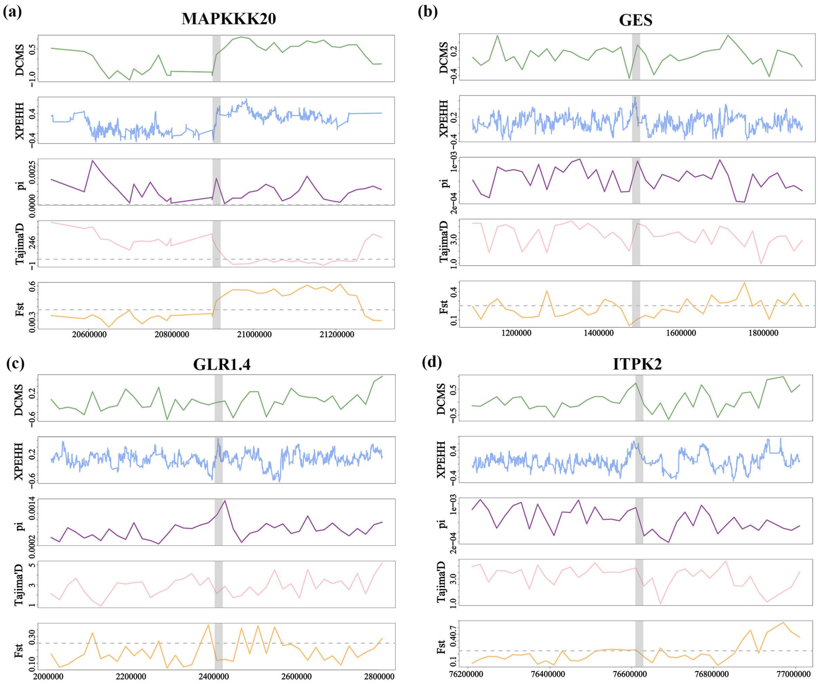


**Figure S16.** Multinumerical line graphs of four key adaptive genes in Danxia landform populations.

A zoom-in on three genetic statistic scores (pi, Tajima's *D*, and $F_{ST}$), XP-EHH values, and DCMS scores, for the *MAPKKK20*, *GES*, *GLR1.4*, and *ITPK2* genes regions and their upstream and downstream 400 kb extension. Each genetic statistic is based on a sliding window analysis using nonoverlapping 20 kb windows.

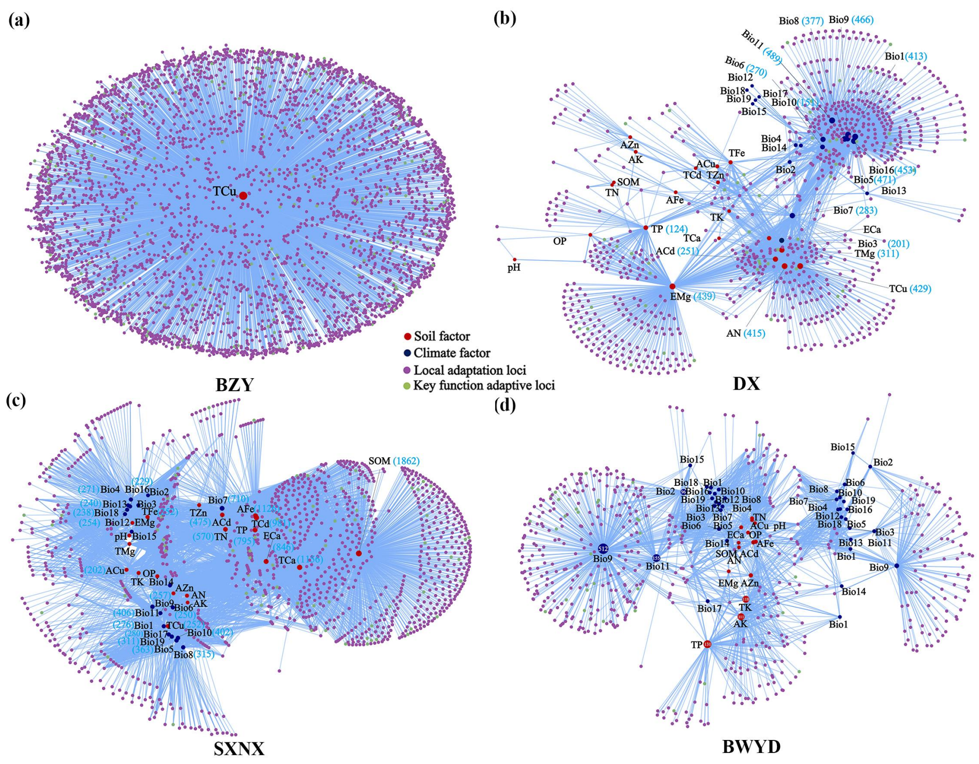


**Figure S17.** Network relationships between environmental variables and local adaptive variants across lineages of *Firmiana danxiaensis.*

Numbers in circles indicate variants significantly influenced by each environmental variable.

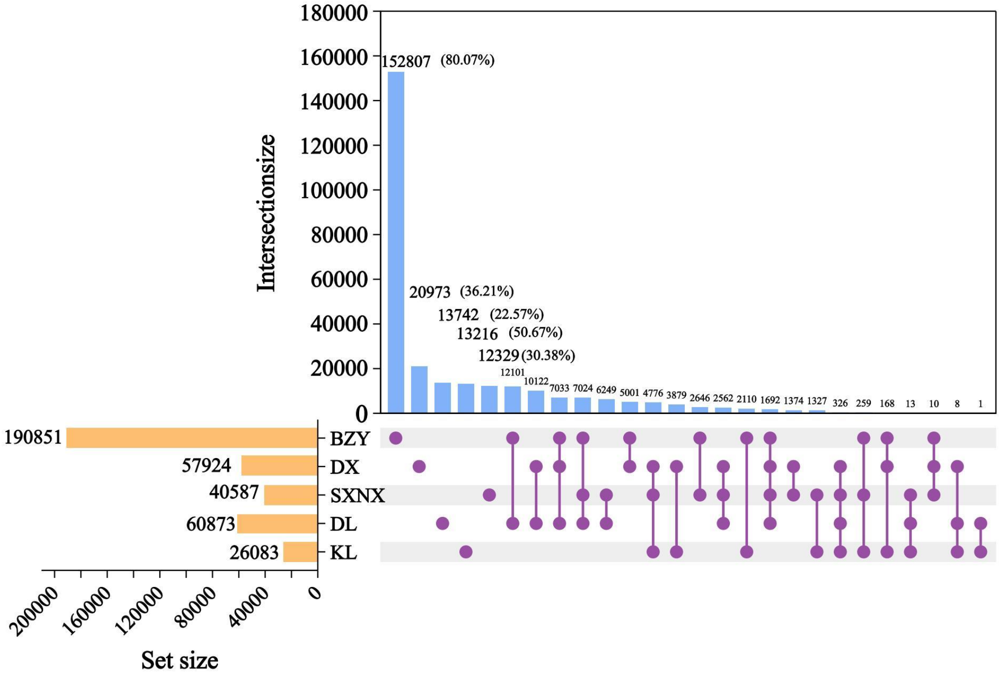


**Figure S18.** Upset plot of positively selected variants across landforms and lineages of *Firmiana danxiaensis.*

Orange bars = total number of selective sweep variants per group; blue bars = overlapping variants across group combinations (purple circles).
